# Metabolomics of healthy and stony coral tissue loss disease affected *Montastraea cavernosa* corals

**DOI:** 10.1101/2021.08.18.456681

**Authors:** Jessica M Deutsch, Olakunle Jaiyesimi, Kelly Pitts, Jay Houk, Blake Ushijima, Brian K. Walker, Valerie J. Paul, Neha Garg

**Author notes:** **Correspondence:** Neha Garg.

## Abstract

Stony coral tissue loss disease, first observed in Florida in 2014, has now spread along the entire Florida Reef Tract and on reefs in many Caribbean countries. The disease affects a variety of coral species with differential outcomes, and in many instances results in whole-colony mortality. We employed untargeted metabolomic profiling of *Montastraea cavernosa* corals affected by stony coral tissue loss disease to identify metabolic markers of disease. Herein, extracts from apparently healthy, diseased, and recovered corals, *Montastraea cavernosa*, collected at a reef site near Ft. Lauderdale, Florida were subjected to liquid-chromatography mass spectrometry-based metabolomics. Unsupervised principal component analysis reveals wide variation in metabolomic profiles of healthy corals of the same species, which differ from diseased corals. Using a combination of supervised and unsupervised data analyses tools, we describe metabolite features that explain variation between the apparently healthy corals, between diseased corals, and between the healthy and the diseased corals. By employing a culture-based approach, we assign sources of a subset of these molecules to the endosymbiotic dinoflagellates, Symbiodiniaceae. Specifically, we identify various endosymbiont-specific lipid classes, such as betaine lipids, glycolipids, and tocopherols, which differentiate samples taken from apparently healthy corals and diseased corals. Given the variation observed in metabolite fingerprints of corals, our data suggests that metabolomics is a viable approach to link metabolite profiles of different coral species with their susceptibility and resilience to numerous coral diseases spreading through reefs worldwide.

## Introduction

Corals are holobionts consisting of the coral animal, photosynthetic dinoflagellate endosymbionts (family Symbiodiniaceae) commonly referred to as zooxanthellae, and a complex microbiome (Rohwer et al., 2002; Maire et al., 2021). Collectively, members of the holobiont contribute to the fitness of the coral animal (Rohwer et al., 2002; Mieog et al., 2009; Gordon and Leggat, 2010; Parkinson et al., 2015). Coral reefs provide a variety of important services, including hosting diverse marine ecosystems (Plaisance et al., 2011; Fisher et al., 2015), supplying pharmacophores for drug development (Bruckner, 2002; Sang et al., 2019), protecting shorelines (Reguero et al., 2021), and contributing to coastal economies with an estimated collective worth of $3.4 billion in the US (Brander and Van Beukering, 2013). Factors including increased ocean temperatures, ocean acidification, pollution, and disease outbreaks present continual challenges to the health of corals (Richardson, 1998; Rosenberg and Ben-Haim, 2002; Rogers and Weil, 2010; Pandolfi et al., 2011; Hoegh-Guldberg et al., 2017; Grottoli et al., 2018; Montilla et al., 2019; Howells et al., 2020). A combination of environmental stressors can increase the vulnerability of corals to disease outbreaks (Maynard et al., 2015; Benkwitt et al., 2020). Coral diseases are notoriously difficult to characterize and the pathogenic agents usually remain unknown (Mera and Bourne, 2018; Vega Thurber et al., 2020). One of the challenges with pathogen-identification is the inherent within-species (intra) and between-species (inter) variability to disease susceptibility. Understanding intraspecies and interspecies variability in genotypes, metabolomes, microbiomes, and other physiological factors such as bleaching history of field corals is critical to investigating their contribution to disease resistance and susceptibility, which are essential to effective management and restoration efforts.

Caribbean coral reefs are experiencing an ongoing outbreak of a tissue loss disease named stony coral tissue loss disease (SCTLD), which first appeared along the Florida Reef Tract in 2014 (Miller et al., 2016; Walton et al., 2018; Muller et al., 2020) and continues to spread throughout various countries in the Caribbean (Precht et al., 2016; Alvarez-Filip et al., 2019; Kramer et al., 2020; Muller et al., 2020; Estrada-Saldívar et al., 2021; Heres et al., 2021; Thome et al., 2021). At least 24 different coral species are susceptible to SCTLD, however, intra- and interspecific differences have been observed (Precht et al., 2016; Aeby et al., 2019; Ushijima et al., 2020; Walker et al., 2020; Meiling et al., 2021). Reefs affected by SCTLD can have coral community composition altered within a few months of initial observation (Walker, 2018; Estrada-Saldívar et al., 2020), highlighting the imminent threat of this disease. While the etiological agent remains unknown (Landsberg et al., 2020; Neely et al., 2020), host-endosymbiont harmony seems to be disrupted (Aeby et al., 2019; Landsberg et al., 2020), disease progression is suggested to be mediated by bacteria (Aeby et al., 2019), and treatment with antibiotics is demonstrated to halt lesion progression for a majority of corals in Florida (Neely et al., 2020; Shilling et al., 2021; Walker et al., 2021). Histological studies indicate some characteristic features of SCTLD (Landsberg et al., 2020); however, without knowledge of the etiological agent there is currently no feasible diagnostic test to confirm SCTLD in diseased corals.

The emergence of -omics studies offers a unique opportunity to elucidate the complex relationships within the members of the coral holobiont at the genetic and chemical level (Joyce and Palsson, 2006; Gordon and Leggat, 2010; Ramos-Silva et al., 2013; Ainsworth et al., 2015; Dixon et al., 2015; Kenkel and Matz, 2016; Drake et al., 2018). Through microbiome sequencing, recent studies described shifts in corals’ microbial community composition upon SCLTD infection (Meyer et al., 2019; Rosales et al., 2020; Becker et al., 2021). However, the metabolomes, the collection of small molecules called metabolites, of these holobionts have not been evaluated. Metabolomics-based approaches provide insight into an organism’s physiological and biochemical state at the time of sample collection (Goodacre, 2007; Viant, 2008; Patti et al., 2012; Wishart, 2019). Thus, metabolomics profiling has been employed to study normal physiology and pathophysiology of diseases. When studying corals, this endeavor is complicated because metabolomic profiles are influenced by the host genotype, the endosymbiont genotype (Symbiodiniaceae), the associated microbiome, the environment, previous history of bleaching, and disease (Muscatine, 1990; Baker, 2003; Roth, 2014; LaJeunesse et al., 2018). Recent application of metabolomics studies on coral holobionts and their environments have demonstrated the applicability of this approach (Sogin et al., 2014; Ochsenkühn et al., 2018; Lohr et al., 2019; Matthews et al., 2020; Roach et al., 2020; Sikorskaya and Imbs, 2020). Direct metabolomics of symbionts have provided insights into their function in the holobiont (Gordon and Leggat, 2010; Hillyer et al., 2017; Sogin et al., 2017; Rosset et al., 2019; Sikorskaya et al., 2021).

Metabolomics studies conducted to date have enabled exploration of changes in metabolite profiles of coral holobionts to specific stressors including thermal bleaching (Quinn et al., 2016b; Sogin et al., 2016; Hillyer et al., 2017; Roach et al., 2021; Williams et al., 2021), but coral metabolomics largely remains an underdeveloped field (Sogin et al., 2014). Two analytical techniques commonly utilized in coral metabolomics include nuclear magnetic resonance (Andersson et al., 2019) and liquid chromatography (LC) coupled to mass spectrometry (MS) (Gordon et al., 2013). Due to the high sensitivity of LC-MS-based approaches and the resulting increased breadth of detected metabolites, LC-MS is a particularly attractive approach towards characterization of metabolome profiles (Lohr et al., 2019). As threats to coral reefs grow, it is important to understand differences in intraspecific chemotypes and genotypes as well as the factors driving these differences. Metabolome studies enhance the understanding of pathogenesis, illuminate the chemical relationships between the coral host, endosymbionts, and other associated microbes, and aid in discovery of biomarkers. There are currently no published studies that compare healthy and SCTLD-infected coral metabolomes or describe the intraspecies variability. Hence, we employed untargeted LC-MS metabolomics to reveal differences in the metabolomic profiles of apparently healthy *Montastraea cavernosa* corals as well as differences between apparently healthy coral colonies and colonies with disease lesions from tissue samples collected at an endemic site in Broward County, FL. Additionally, we used a culture-based approach to co-analyze the metabolomes of three Symbiodiniaceae species to reveal that at least a third of the top metabolites separating the field coral sample groups belonged to Symbiodiniaceae. We identify a number of Symbiodiniaceae metabolites using a suite of recently developed metabolite feature annotation tools. This study highlights that the culture-based approach, when combined with metabolomics, facilitates an understanding of the contribution of different holobiont components to the metabolomic profiles, which can aid in teasing apart the source of metabolites that dictate the separation in metabolomic profiles. We propose that in addition to the prominent contribution of Symbiodiniaceae to metabolomes of field corals, careful cataloging of host genotype and past environmental insults is important to understand intraspecies differences of field corals.

## Materials and Methods

### Sample collection, data collection and analyses workflow

#### Site selection

An experimental site was set up on March 23, 2020 located 475 m from shore (26° 9.05’ N, 80° 5.75’ W), on the Broward-Miami Ecoregion Ridge-Shallow habitat (Walker et al., 2008; Walker, 2012) in about 7 m depth (Figure 1). Nineteen visibly diseased *Montastraea cavernosa* colonies were tagged, photographed, and mapped across a 100m x 35m area of hardbottom using a floating GPS to obtain coordinates for all colonies. Healthy-looking colonies were not mapped. Collections were delayed due to the coronavirus pandemic. This site was revisited August 19, 2020 to evaluate disease progression and sample corals. At this time, 6 newly diseased colonies were additionally sampled and three previously diseased colonies no longer showed signs of disease (apparently recovered). Thus, 22 diseased colonies and 3 recovered colonies were sampled in August. On October 14, 2020, three new diseased colonies and ten colonies completely covered in seemingly healthy tissue were added to the site and sampled for metabolomics.

**Figure 1.**
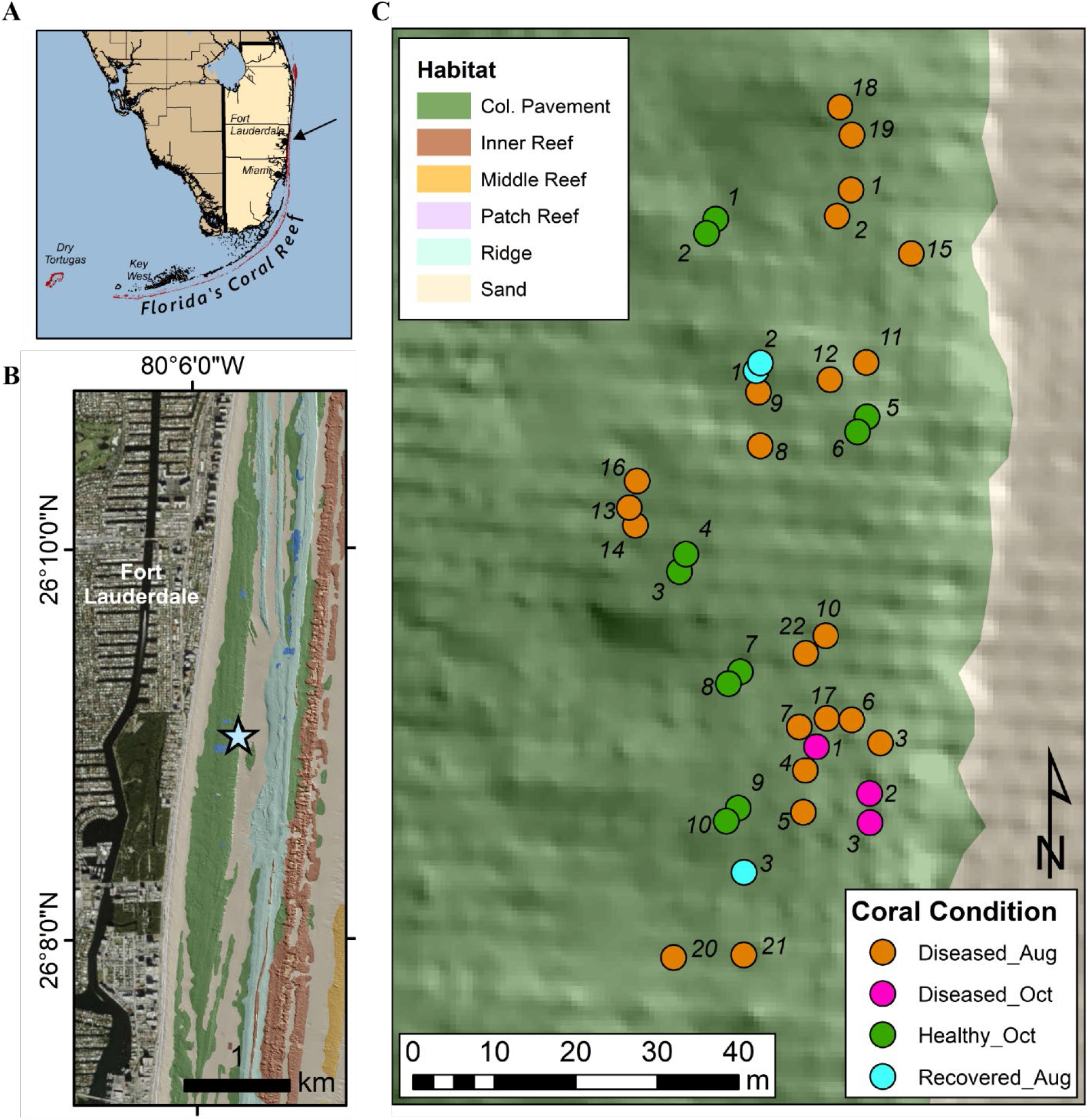
Site map of sample collection site in Broward County, FL. (A) Panel A shows the southern coast of FL with an arrow indicating the sampling location. (B) Panel B shows the sample site (star) in relation to local benthic habitats and topography as described in Walker et al. (2008). (C). Panel C describes the location of the sampled corals, as well as their health condition at the time of sampling.

#### Sampling and extraction procedure for coral samples

The samples were collected by divers on SCUBA with a 10 mL needleless syringe by agitating 4-5 adjacent polyps with the tip of the syringe. Dislodged tissue fragments and mucus were slowly drawn into the syringe until 10 mL of sample was collected. Diseased colonies were sampled along the disease lesion. Recovered colonies that displayed signs of SCTLD in March 2020, but no longer appeared to be diseased were sampled at the furthest point away from the former disease lesion to sample an area that was least likely to be impacted by SCTLD. For healthy corals, colonies that were completely covered in apparently healthy tissue were chosen to ensure that they were less likely to have had SCTLD or other tissue loss. All samples were then placed in 15 mL conical tubes, frozen and lyophilized.

After weighing, the lyophilized samples were transferred to a 20 mL scintillation vial for extraction. To each conical tube, 6 mL of 2:1 methanol (MeOH): H_2_O was added to dissolve any residual material left in the tube and transferred to the scintillation vial containing the lyophilized sample. To this vial, 4 mL of ethyl acetate (EtOAc) was added, and the samples were extracted (10 ml of 2:2:1 EtOAc: MeOH: H_2_O) for 4 h at 23 °C. The extracts were spun in a Savant SpeedVac for 10 min, and the supernatant was transferred to a 7 mL glass vial (precleaned three times with MEOH). The solvent was removed *in vacuo* using the Savant SpeedVac set at 35 °C and then lyophilized at −45 °C to remove any remaining water. The concentrate was transferred back into the initial falcon tube in 4 mL of 3:1 MeOH: H_2_O and centrifuged at 4500×g for 10 min to remove particulates. For metabolomics, 0.5 mL of the supernatant was transferred to a pre-weighed 1.5 mL microcentrifuge tube, centrifuged, and lyophilized. The remaining supernatant was transferred back to the 7 mL scintillation vial and dried *in vacuo*. The samples were stored at −20 °C until LC-MS data was acquired.

#### Sampling and extraction procedure for cultured endosymbionts

Symbiodiniaceae isolates, *Breviolum, Symbiodinium*, and *Durusdinium*, from the University of Buffalo Undersea Reef Research (BURR) collection were received from Richard Karp and Andrew Baker, University of Miami. These endosymbionts were isolated by Mary Alice Coffroth from *Orbicella faveolata* corals collected in the Florida Keys between 2002 and 2005. All isolates were cultured in F/2 media, incubated at 27 °C, with 20 μE of light on a 14:10 diurnal cycle. To prepare extracts, 500 μl of each culture was transferred to a 1.5 mL microcentrifuge tube. Cells were pelletized at 500×g for 5 min to remove media and then washed twice with seawater filtered through a 0.22 μm-pore membrane filter (FSW). A sample of 0.2 μm-filtered seawater (500 μL) was extracted in tandem as a control. The washed pellet was resuspended in 100 μL of HPLC grade water for transfer into a pre-cleaned 7 mL scintillation vial and then frozen and lyophilized. Samples were extracted overnight at −20 °C in 4 mL of 2:2:1 EtOAc: MeOH:H_2_O. Solvents were removed on a SpeedVac at 35 °C for 4 h. The dried samples were resuspended in 3:1 MeOH:H_2_O and transferred into a 1.5 mL microcentrifuge tube. The samples were centrifuged at 4500×g for 10 min, and the supernatant was transferred to the pre-weighed microcentrifuge tube. This supernatant was removed via SpeedVac for 3 h after which the concentrated extract was frozen, then lyophilized. The weight of lyophilized extract was recorded.

#### Mass Spectrometry Data Acquisition and analyses

All dried extracts were resuspended in 100% MeOH (LC-MS grade) containing 1 μM of sulfadimethoxine as an internal standard. The samples were analyzed with an Agilent 1290 Infinity II UHPLC system (Agilent Technologies) using a Kinetex 1.7 μm C18 reversed phase UHPLC column (50×2.1 mm) coupled to an ImpactII ultrahigh resolution Qq-TOF mass spectrometer (Bruker Daltonics, GmbH, Bremen, Germany) equipped with an ESI source for MS/MS analysis. MS spectra were acquired in positive ionization mode, *m/z* 50–2000 Da. An active exclusion of two spectra was used, implying that an MS^1^ ion would not be selected for fragmentation after two consecutive MS^2^ spectra had been recorded for it in a 0.5 min time window. The exclusion was reconsidered and additional MS^2^ spectra was acquired if five-fold enhancement in intensity was observed. The eight most intense ions per MS^1^ spectra were selected for further acquisition of MS^2^ data. The chromatography solvent A: H_2_O + 0.1% v/v formic acid and solvent B: MeCN + 0.1% v/v formic acid were employed for separation. Flow rate was held constant at 0.5 mL/min throughout the run. The gradient applied for chromatographic separation was 5% solvent B and 95% solvent A for 3 min, a linear gradient of 5% B–95% B over 17 min, held at 95% B for 3 min, 95% B–5% B in 1 min, and held at 5% B for 1 min, 5%B-95%B in 1 min, held at 95% B for 2 min, 95%B-5%B in 1 min, then held at 5%B for 2.5 min at a flow rate of 0.5 mL/min throughout. A mixture of 6-compounds (amitryptiline, sulfamethazine, sulfamethizole, sulfachloropyridazine, sulfadimethoxine, coumarin-314) was run as quality control every 8 samples to ensure consistent instrument and column performance.

The raw data was converted to mzXML format using vendor software and preprocessed on MZmine 2.53 using mass detection, chromatogram building, chromatogram deconvolution, isotopic grouping, retention time alignment, duplicate removal, and missing peak filling (Pluskal et al., 2010). This processed data was submitted to the feature-based molecular networking workflow on the Global Natural Product Social Molecular Networking (GNPS) platform. Herein, the output of MZmine includes information about LC-MS features across all samples containing the *m/z* value of each feature, retention time of each feature, the area under the peak for the corresponding chromatogram of each feature, and a unique identifier. The MS^2^ spectral summary contains a list of MS^2^ spectra, with one representative MS^2^ spectrum per feature. The mapping information between the feature quantification table and the MS^2^ spectral summary was stored in the output using the unique feature identifier and scan number, respectively. This information was used to relate LC-MS feature information to the molecular network nodes. The quantification table and the linked MS^2^ spectra were exported using the GNPS export module (Pluskal et al., 2010; Nothias et al., 2020) and the SIRIUS 4.0 export module (Pluskal et al., 2010; Dührkop et al., 2019). Feature Based Molecular Networking was performed using the MS^2^ spectra (mgf file) and the quantification table (csv file). Three quantification tables were generated; one with LC-MS feature list of data acquired on extracts of all coral samples analyzed in the study, the second with LC-MS feature list of data acquired on the extracts of all coral samples and extracts of cultured Symbiodiniaceae, and the third with LC-MS feature list of data acquired on the extracts of coral samples collected in October and extracts of cultured Symbiodiniaceae.

Two molecular networks were generated using this workflow. To generate each network, the data was filtered by removing all MS/MS fragment ions within +/- 17 Da of the precursor *m/z*. MS/MS spectra were filtered by choosing only the top 6 fragment ions in the +/- 50 Da window throughout the spectrum. The precursor ion mass tolerance was set to 0.02 Da and the MS/MS fragment ion tolerance to 0.02 Da. A molecular network was then created where edges were filtered to have a cosine score above 0.7 and more than 4 matched peaks. Further, edges between two nodes were kept in the network if and only if each of the nodes appeared in each other’s respective top 10 most similar nodes. Finally, the maximum size of a molecular family was set to 100, and the lowest scoring edges were removed from molecular families until the molecular family size was below this threshold. The spectra in the network were then searched against GNPS spectral libraries (Wang et al., 2016). The library spectra were filtered in the same manner as the input data. All matches kept between network spectra and library spectra were required to have a score above 0.7 and at least 4 matched peaks. The molecular networks were visualized using Cytoscape version 3.7.2 (Shannon et al., 2003). The compound annotations follow the “level 2” annotation standard based upon spectral similarity with public spectral libraries or spectra published in the literature as proposed by the Metabolomics Society Standard Initiative (Sumner et al., 2007). Annotations were confirmed with commercial standards when available. The MS2LDA analysis was performed as previously described with default parameters (van der Hooft et al., 2017; McAvoy et al., 2020). The output of this analysis is available at https://gnps.ucsd.edu/ProteoSAFe/status.jsp?task=a7dfdc1606c148c1bbda9c0fbc803d2e.

Prior to statistical analysis, blank subtraction was performed on the quantification tables using a Juptyer notebook, available at [GitHub-Garg-Lab/Jupyter-Notebook-Blank-Subtraction-Broward-County-Corals](https://github.com/Garg-Lab/Jupyter-Notebook-Blank-Subtraction-Broward-County-Corals). The mean of each feature in the samples is compared separately to the mean of the feature in the solvent controls and LC-MS blanks. A feature is retained if its sample mean is greater than 0.25 × the mean of the sample in the blanks. The entire quantification table is exported, with each feature marked as “true” (feature is retained) or “false” (feature is not retained). Visualization of the molecular network, metadata mapping, and feature filtering was performed using Cytoscape (v 3.7.2) (Shannon et al., 2003). Within Cytoscape, node filtering was performed using metadata and quantification tables and the nodes corresponding to features present in blank were removed. All Euler diagrams were created using EulerAPE (v 3.0.0) (Micallef and Rodgers, 2014).The filtered quantification table were submitted to MetaboAnalyst 5.0 for all multivariate analysis (Chong et al., 2019). Pareto scaling was used for normalization, then principal component analysis and hierarchical clustering using the Euclidean distance measure and Ward algorithm for clustering were performed. Heat maps of discriminating features ranked using non-parametric ANOVA were generated (Supplementary file 1). The software PRIMER version 7 was used to test similarities in metabolite species between sampled corals with differing health conditions. Cluster analyses with SIMPROF tests and non-metric multi-dimensional scaling (MDS) plots were constructed from a Bray-Curtis similarity matrix of log transformed metabolite data. The metabolite similarities between corals using the coral condition status at the time of sampling were tested using analyses of similarities (ANOISM). The similarity percentages (SIMPER) were calculated for each cluster to identify the metabolites most responsible for the differences between those clusters with 94% similarity.

Additional compound annotations were performed using the XCMS Online, and *in silico* tools such as MolDiscovery (v. 1.0.0) and SIRIUS 4.5 with CSI:FingerID and CANOPUS, and through literature searches for compounds known to be produced by corals and their endosymbionts for untargeted analysis. MolDiscovery, available through the GNPS platform, compares *in silico* generated mass spectra of small molecules from a variety of databases with experimental MS^2^ spectra and includes a similarity score for each reported match (Cao et al., 2020). The LC-MS feature list from the extracts of coral samples collected in October (mgf file) was submitted to the MolDiscovery workflow. The mzXML files of data acquired on extracts of corals collected in October were submitted to the XCMS Online platform and processed using the parameters for UPLC Bruker Q-TOF pos (6675) (Gowda et al., 2014).

CSI:FingerID was employed using default settings. The fragmentation trees were computed for submitted LC-MS features, and predicted chemical formulas and structures that best matched the experimental spectra were ranked using machine learning (Dührkop et al., 2019). The compound classes were assigned using CANOPUS, which uses MS/MS spectra and the chemical structures predicted by CSI:FingerID to propose chemical classes for a feature (Djoumbou et al., 2016; Dührkop et al., 2021).

## Results and Discussion

### Metabolomic profiles of all samples

The apparently healthy and diseased *M. cavernosa* samples were collected in two batches. The first batch was collected in August of 2020 (samples labelled as Diseased_Aug, 22 samples) and the second batch was collected in October of 2020 (samples labelled as Diseased_Oct, 3 samples and Healthy_Oct, 10 samples) (Figure 1 and Supplementary Figure 1). Additionally, three samples that were previously observed to have signs of SCTLD when the site was set up in March 2020, but no longer displayed signs of disease were also collected from the same site (samples labeled as Recovered_Aug). The extracted metabolites were stored frozen at −20 °C in a dried form and then the metabolomics data acquisition was performed in a single run to avoid batch-to-batch variation. The area under the curve for each detected MS^1^ feature was obtained from extracted ion chromatograms using MZmine2. The resulting feature table was analyzed with a suite of data visualization and annotation tools, which are described below and detailed in the methods (Figure 2). After removing metabolite features detected in solvent controls, 3914 metabolite features were detected in the coral extracts. The corresponding data on these features was first subjected to principal component analysis (PCA) to visualize metabolomic variation between samples. When PCA was performed on all samples, 44.3% of the total variation was explained by the PCA components 1 and 2 (Figure 3A). This PCA indicated relatively separate clustering of the healthy (n=10, green dots) from the diseased samples (n=22, pink and orange dots), however the recovered samples (n=3, cyan dots) did not all cluster with healthy or diseased samples and showed large within-group variation similar to the diseased samples. The first PCA component explained the variation within each sample class, whereas the second component explained separation between the healthy and diseased corals. A permutational multivariate analysis of variance (PERMANOVA) was performed with 999 permutations to investigate if the metabolome of diseased and healthy stony corals differed significantly. The PERMANOVA *F*-test with F (p) value of 4.834 (0.001) supported the observation that the metabolomes differ between healthy and diseased coral samples. Next, we employed partial least square discriminant analysis (PLS-DA), which as opposed to PCA, is a supervised method and maximizes the separation between pre-defined groups. The PLS-DA model revealed clear separation between healthy and diseased samples, while still highlighting within-group variation with lower Q^2^ value of 0.64 (Supplementary Figure 2). Such within-group variation was previously shown for white syndrome disease in *Acropora* and *Playtgyra* (Ochsenkühn et al., 2018). Thus, within-group variation is a hallmark of metabolomic profiles of coral holobionts collected in the wild and should be carefully considered in study design employing –omics methods. The metabolomics variation among healthy corals in a nursery has also been investigated for at least one species of coral and was attributed partly to the host genotype, while other factors were proposed for with-in group variation (Lohr et al., 2019). The metabolome profile variation in visually healthy corals in the ocean may be attributed to multiple sources, including host genotype, associated endosymbionts (Symbiodiniaceae), the microbiomes, the surrounding abiotic environment, past bleaching history, and whether these corals are under stress before visual signs of disease appear. The variation in the diseased samples may additionally arise from severity of disease, duration of infection, or variation in groups of colonizing microorganisms.

**Figure 2.**
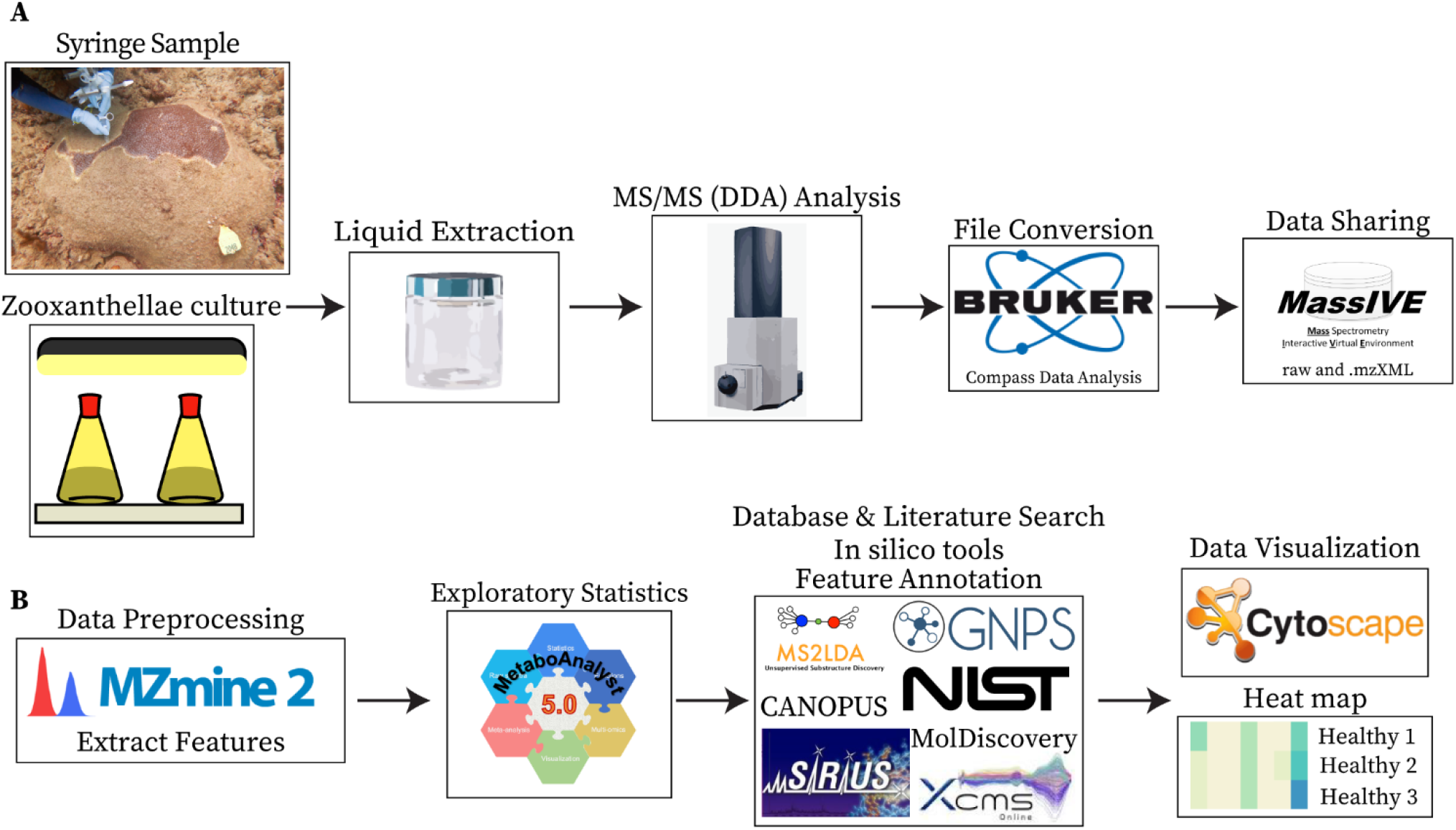
Sample collection, data collection, and analyses. **(A)** Panel A shows the sample types analyzed (syringe samples collected from coral colonies and cultured Symbiodiniaceae) and the steps employed for data acquisition, conversion, and sharing. **(B)** Panel B shows the steps employed for data processing, visualization, and the tools utilized for compound annotations.

**Figure 3.**
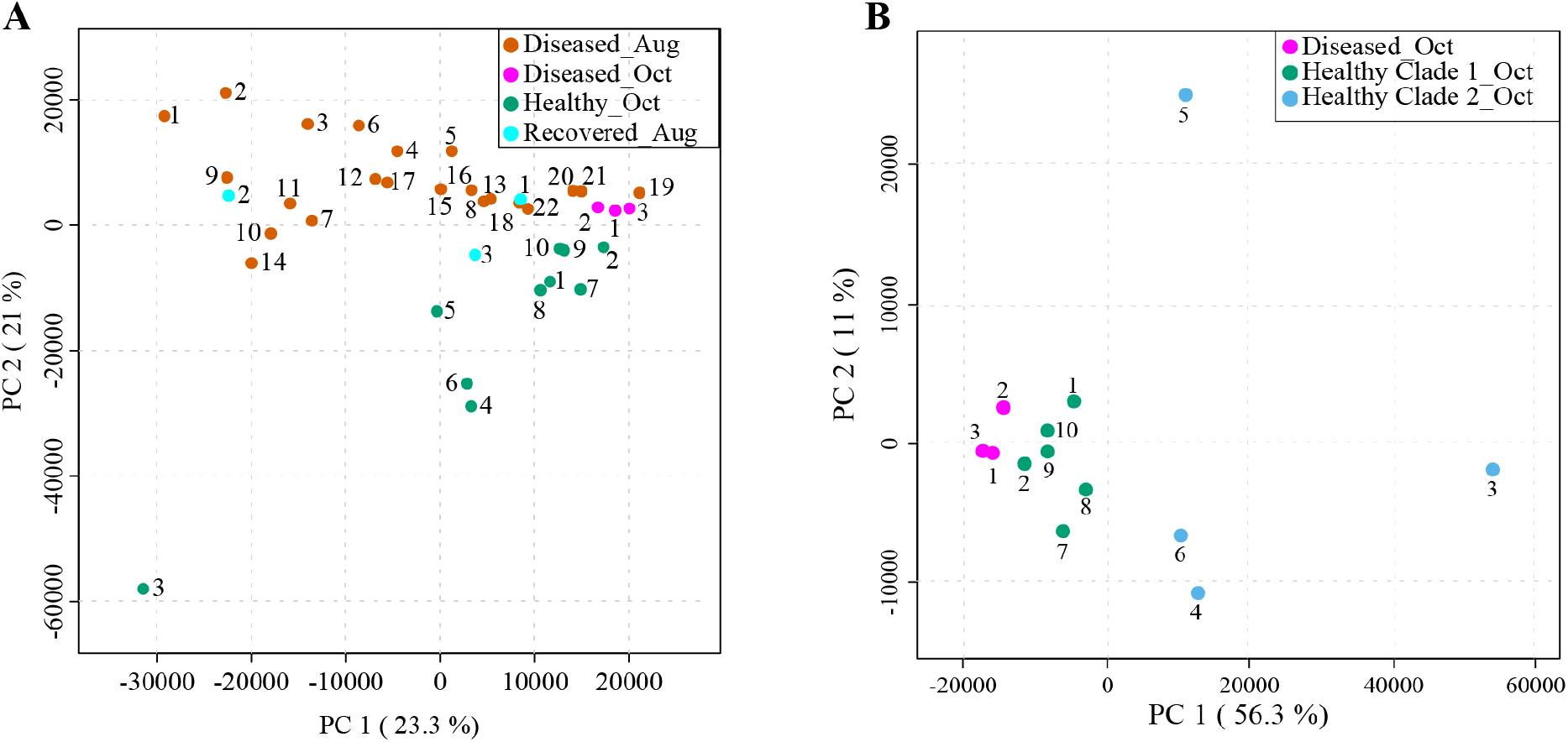
Principal component analyses (PCA) of LC-MS-based metabolomics data acquired on syringe samples from *M. cavernosa* corals in Broward county, FL. **(A)** Panel A shows PCA of all samples collected in August and October: Diseased_Aug (orange), Diseased_Oct (pink), Healthy_Oct (green), and Recovered_Aug (cyan). The healthy samples were collected only in October, whereas diseased samples were collected in both August and October. **(B)** Panel B shows PCA of healthy (green and blue dots), and diseased (pink dots) samples collected in October. The healthy samples are displayed in two colors based on two clades observed for healthy samples in HCA analyses shown in Figure 4 and Supplementary Figure 3. The amount of variance explained by each principal component is shown in parentheses on the corresponding axis.

### Metabolomic profiles of samples collected in October

Assessing the metabolomes of apparently healthy and diseased corals collected in October alone revealed tighter clustering of the diseased samples with relatively separate clustering from healthy samples suggesting that the disease progression may lead to convergence of metabolomes (pink dots, Figure 3B). The clustering of the diseased samples collected in October was tighter than the August diseased samples (Figure 3A). In this regard, the three principal components for October samples explained 77% of the total variation with component 1 accounting for the majority of the variation (56.3%) as compared to 23.3% of the variation explained by PC1 when all samples were considered (Figure 3 and Supplementary Figure 3A). Furthermore, the healthy samples 3-6 displayed greater spread across all components suggesting large within-group variation in these apparently healthy samples. Among these, the healthy 3 showed the largest variation along all three components in PCA and displayed grey pigmentation of the entire colony (Supplementary Figure 1). Thus, the phenotypic variation between the coral colonies of the same species in the field is captured at the metabolome level suggesting metabolomics approaches could be productive in the future for linking specific phenotypes to chemotypes as databases for compounds detected in coral holobionts are populated. The *M. cavernosa* corals have been suggested to maintain high levels of genetic diversity compared to other coral species (Budd et al., 2012). The relative intraspecies variation in metabolomic profiles of different species of corals in the future will enable insights into whether genetic diversity can be a driver of observed metabolomic variation.

### Geographic proximity and metabolomic variation

To visualize the relation of spatial proximity of coral colonies to observed clustering in PCA, we categorized the corals by the 94% similarity MDS clusters in a map and circled colonies with close relationships in the hierarchical cluster analysis (HCA) (Figure 4), wherein samples are clustered in a dendrogram in which the linkages and their relative proximity equates to the similarity of metabolomic profile. The closer the branches, the more similar are the given object’s variables, herein metabolite abundances. Two major clusters were observed in HCA (branches 1 and 2, Figure 4A and Supplementary Figure 3B). Each cluster was further subdivided in two sub-clusters (Figure 4A, branches 3, 4 and branches 5, 6). In each of these sub-clusters, all apparently healthy samples clustered together whereas diseased samples separated into additional sub-clusters. The Recovered_Aug samples did not follow a defined pattern; Recovered_1 and Recovered_3 samples clustered with healthy samples whereas the Recovered_2 sample clustered with the diseased samples. To query where this clustering is dictated by the proximity of coral colonies, the individual branches in HCA were colored based on their location shown in the site map (Figure 4A). Indeed, a number of neighboring branches revealed geographic proximity, however, this observation was not consistent across all colonies within the HCA groups. The yellow branches in the HCA and corresponding yellow circles in the map highlight connected colony pairs that were close neighbors. The purple and orange highlighted branches and circles were colonies in the same HCA sub-cluster, but were not directly connected. Some colonies in these branches were also in close spatial proximity. Based on the two clusters observed in HCA (Figure 4A branches 4 and 6), the healthy samples were colored in green and blue in the PCA plot in Figure 3B. The clustering patterns were reproducible for diseased corals when analyses was performed on data generated on diseased samples only (Supplementary Figure 3C).

**Figure 4.**
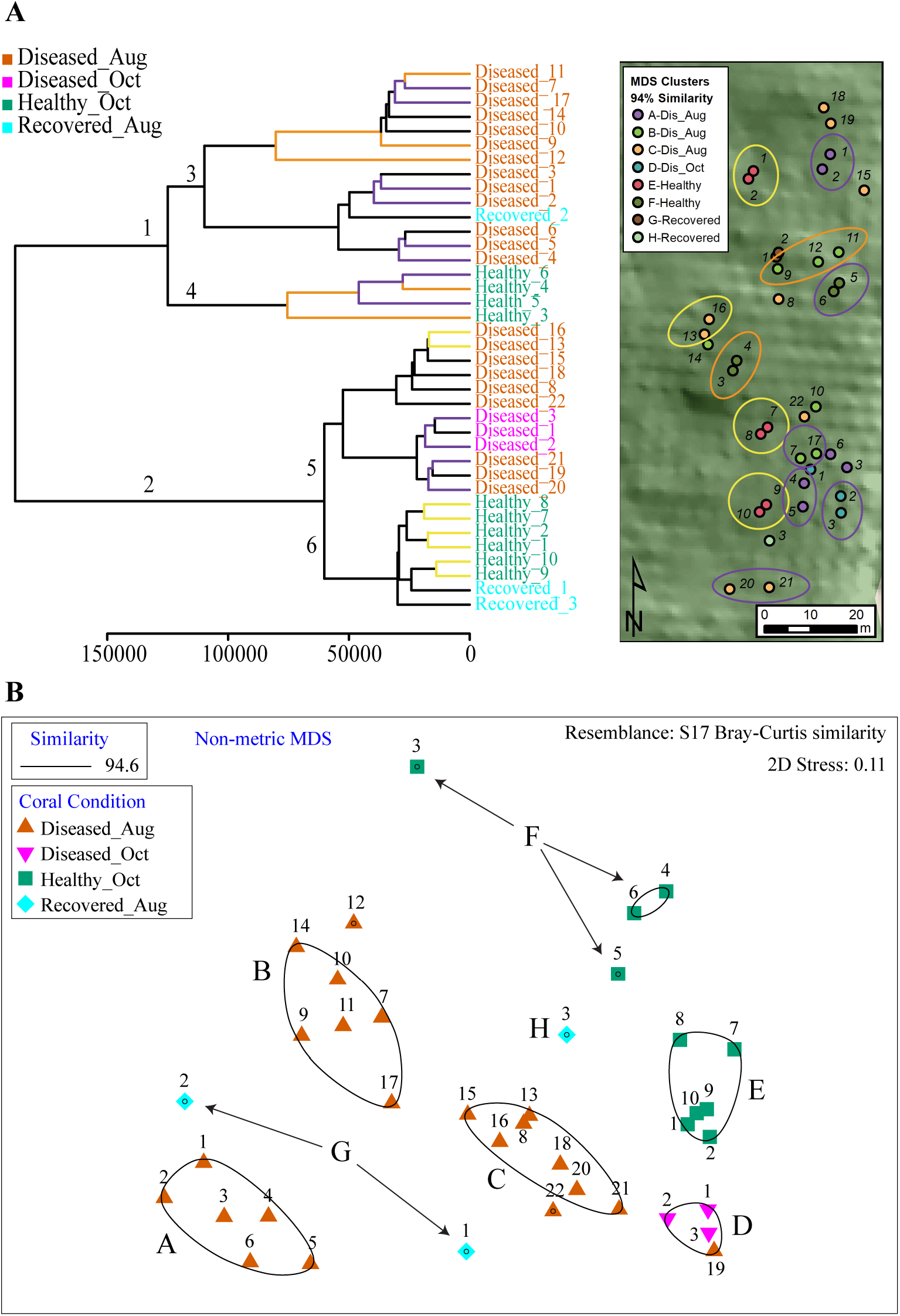
The HCA and MDS plot of all samples analyzed in this study. **(A)** The healthy samples were divided across 2 major clades (clades 4 and 6) and served as basis to color healthy samples in PCA plots as green and blue in Figure 3B. The branches in HCA are colored based on location proximity deciphered from site map. The branches in yellow represent samples that are within the same sub-clade and are neighbors in the site map shown as circles (For e.g., sample healthy 1 and 2 are immediate neighbors in site map). The branches in purple and orange are present across two sub-clades. The branches with either color represent neighboring samples in site map. For e.g., healthy sample 3 and 4 in orange branches are neighbors in site map. Similarly, healthy samples 5 and 6 in purple branches are neighbors in site map. **(B)** Panel B shows output of MDS analysis where diseased samples are grouped in four clusters and healthy samples are grouped in two clusters.

The non-metric multi-dimensional scaling (MDS) plots divided the samples into eight distinct clusters (Figure 4B). Categorizing the corals by the 94% similarity clusters in the map revealed no obvious large-scale spatial clustering. Many colonies with similar metabolite communities were spread throughout the site and spatially comingled with dissimilar colonies. However, some neighboring colony pairs were close in similarity indicating potential within-MDS-cluster spatial patterns. For example, MDS cluster A indicated colonies 1 and 2 were most similar to each other and furthest away from the other colonies in the similarity group, which was also the case in the map. So was the case with cluster E colonies 7 and 8. However there were many more cases where colonies with dissimilar metabolite communities were in close spatial proximity.

Analyses of Similarities (ANOSIM) were used to test for similarities in metabolites between corals using the coral condition status at the time of sampling. Since three distinct clusters were evident at 94.6% similarity in the Diseased_Aug samples, similarity percentages (SIMPER) were calculated for each cluster to identify the metabolites most responsible for the differences between those clusters. A number of features describing the differences between the healthy and diseased samples include metabolites produced by Symbiodiniaceae such as glycolipids, betaine lipids, and tocopherols and are described below.

The variation in coral metabolomes is a hallmark of complex interactions between the holobiont members that differ in genotypes, chemotypes, as well the varying levels of disease progression observed in field samples. The observations from a smaller number of field samples in October in our study require validation in future studies that employ time series design post the onset of SCTLD. Metabolomics paired with description of host and endosymbiont genotype as well as microbiome structure and bleaching history is essential to explain underlying factors resulting in observed clustering. When coupled with metaproteomics and metatranscriptomics, the underlying mechanisms of such variation can be further explained at the biochemical level.

### Description of metabolite features underlying variations in metabolomic profiling

We used feature-based molecular networking, and various other *in silico* approaches listed in Figure 2 and described in the methods sections, to annotate metabolite features showing within group and between group variations. The distribution of metabolite features across healthy, diseased, and recovered samples was visualized as UpSet plot (Figure 5). Of 3914 metabolite features, 2231 features were observed in all coral samples. Among the metabolite features unique to each group, the largest number of unique features were detected in diseased corals collected in August (366 features) followed by apparently healthy samples (234 features), recovered samples (29 features) and the least number of unique features were detected in diseased coral samples collected in October (11 features). An important component to understanding the response of corals to disease is the ability to assign the source of the metabolites; that is whether the metabolites originate from the host, endosymbionts, the associated microbiome, or the environment. We hypothesized that the large proportion of the metabolome unique to healthy samples may be attributed to the endosymbiotic dinoflagellates, Symbiodiniaceae, as these symbionts display signs of stress during disease and some diseased coral colonies, including all three collected in October, did display bleaching along the tissue margin of the lesion while others did not. The *M. cavernosa* has been reported to display bleaching along the margin of the lesion in response to SCTLD (Aeby et al., 2019). To examine metabolite sources, we cultured three genera of Symbiodiniaceae, extracted the metabolites from the cultures, and acquired untargeted metabolomics data as described for coral samples. The cultured Symbiodiniaceae include *Symbiodinium sp., Durusdinium sp*., and *Breviolum sp*. The MS/MS spectra from the coral tissue extracts and the cultured Symbiodiniaceae were preprocessed together using MZmine. Of the 3992 features retained after blank subtraction, 20.3% of these features were shared between the coral holobiont and the cultured symbionts, with 74.3% of the features unique to the coral holobiont, and 5.4% of features unique to cultured Symbiodiniaceae (Figure 6A). The metabolites produced in culture by three species of cultured Symbiodiniaceae were also analyzed and significant overlap in their metabolomes was observed, as well as unique metabolites detected in culture extracts of each species (Figure 6B). To visualize features driving the differences between various samples types in PCA, a heat map was generated on top-50 metabolite features ranked using non-parametric ANOVA (Figure 7). The MS^2^ spectra for these top 50 features that showed differential abundance between healthy, recovered, and diseased samples from the output of ANOVA were searched using GNPS, NIST 2017, MolDiscovery, XCMS Online, and SIRIUS with CSI:FingerID. Where annotations were not possible, the chemical class was assigned using CANOPUS. These database matches, proposed compound class by CANOPUS, and putative molecular formulas assigned by SIRIUS with CSI:FingerID are reported in Supplementary Table 1. Many of the features driving the separation among individuals within the healthy coral category as well as between the healthy and the diseased coral samples were detected in both the cultured Symbiodiniaceae and field corals (Figure 7, indicated by +/-). Even though these Symbiodiniaceae were not isolated from the *M. cavernosa* coral colonies analyzed in the present study (see methods section) and the cultured Symbiodiniaceae can differ phenotypically from when they are present in the coral host (Maruyama and Weis, 2021), the significant overlap observed between field corals and cultures suggests that a culture-based approach is a viable step to tease apart at least a portion of Symbiodiniaceae metabolomic contributions to the coral holobiont. Indeed by manipulating culturing conditions, various different metabolites were shown to be produced by Symbiodiniaceae (Nakamura et al., 1998). Similar approaches have been used to identify microbiome-specific metabolites in infectious diseases in humans (Quinn et al., 2016a; Garg et al., 2017; Melnik et al., 2019). The variability in the Symbiodiniaceae metabolites within healthy samples was apparent from the heat map and may arise from different host genotypes or differences in associated microorganisms. It is now well known that some coral colonies can display resilience to disease and the role of Symbiodiniaceae in this resilience has been proposed (Correa et al., 2009; Littman et al., 2010; Rouzé et al., 2016; Rosset et al., 2019). Thus, in the absence of mass spectral databases for coral symbionts and microbiomes, a culture-based approach can be employed to further assess their role in metabolome variation in field corals. An alternative approach to culturing allowing direct comparison is to acquire metabolomics data on isolated in hospite pure Symbiodiniaceae fractions from the coral tissue (Sikorskaya et al., 2021).

**Figure 5.**
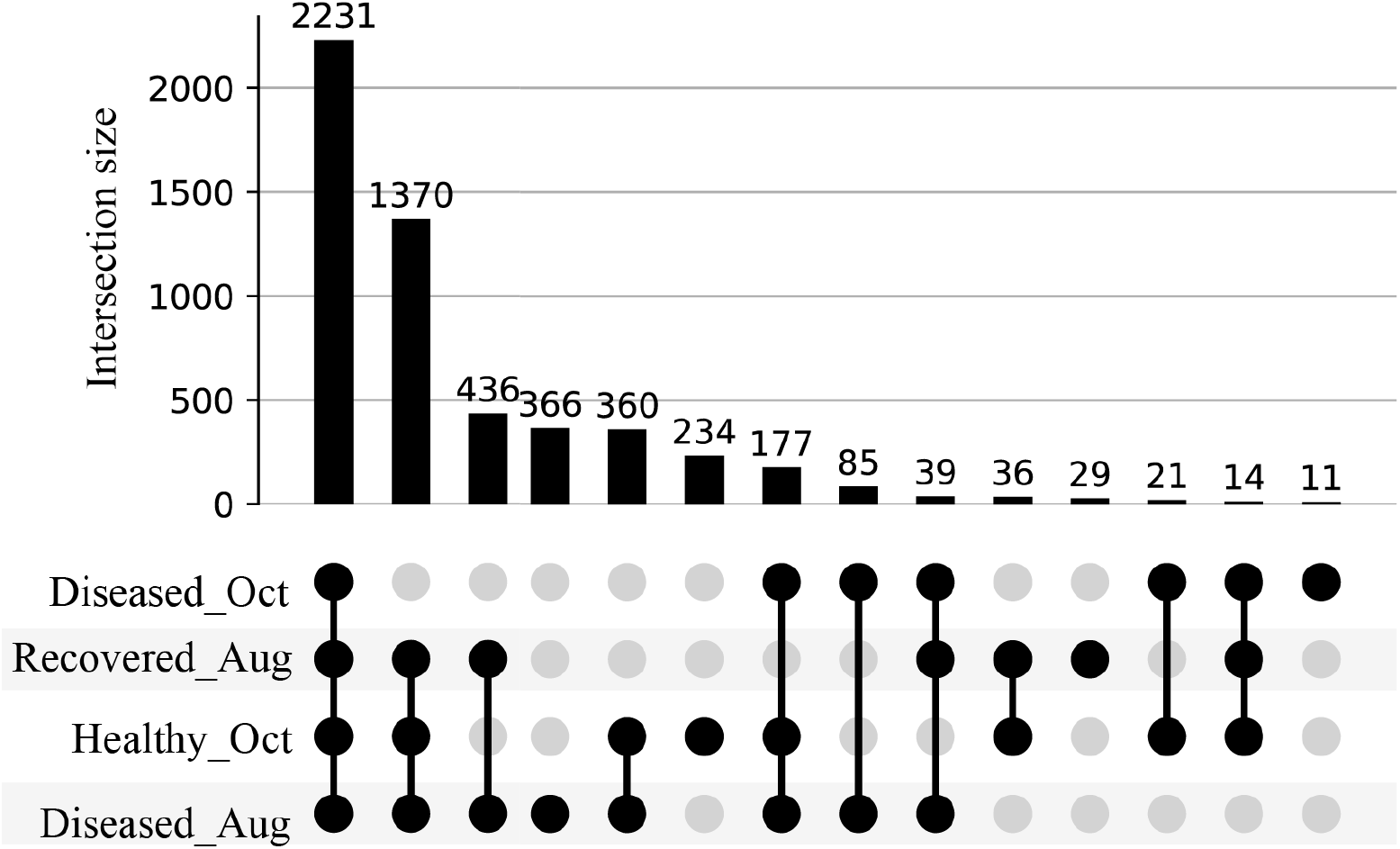
UpSet plot of metabolite features detected in the healthy, diseased, and recovered samples arranged by decreasing frequency. The number on each bar represent total number of metabolite features in the corresponding intersection. For e.g., 2231 represents the detection of a total of 2231 metabolite features across all diseased, healthy, and recovered samples, whereas 1370 represents the detection of a total of 1370 metabolite features across diseased samples collected in August, healthy, and recovered samples.

**Figure 6.**
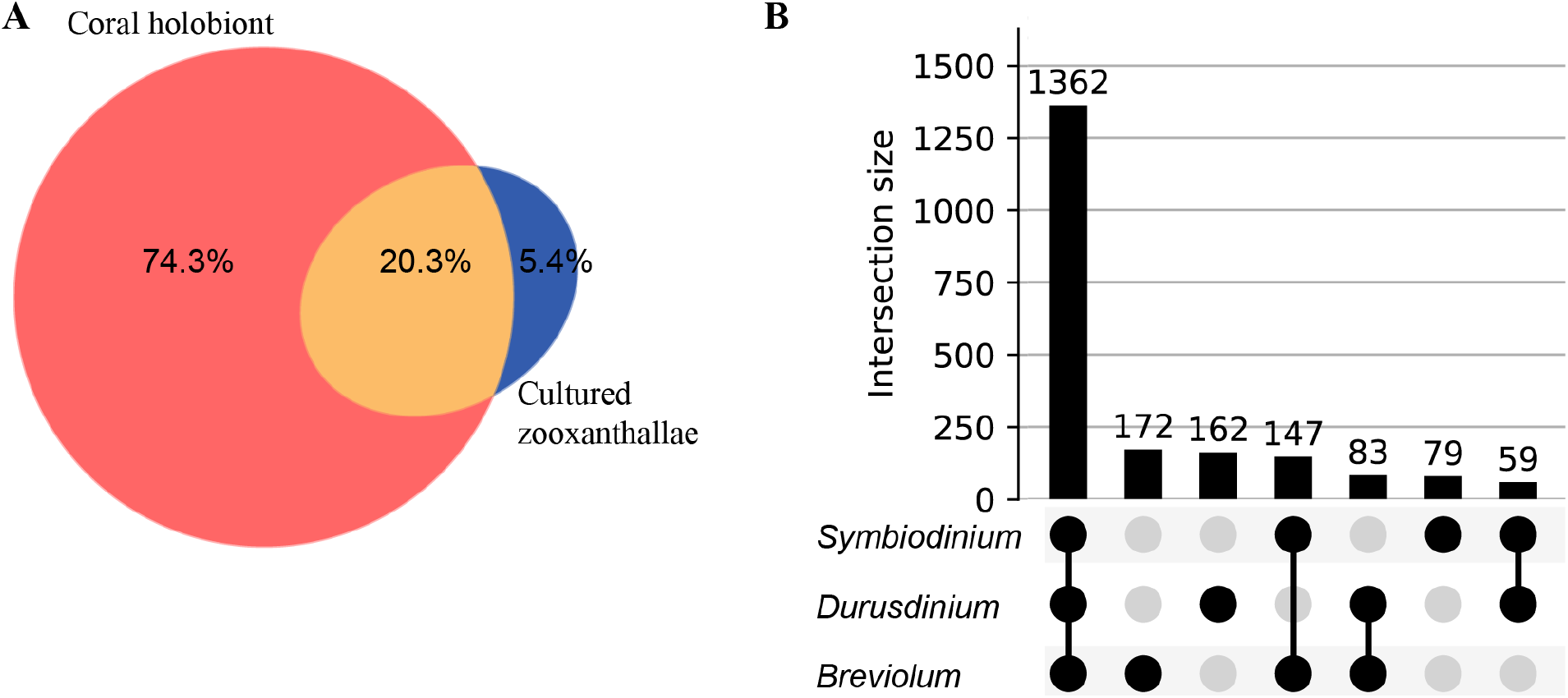
Venn diagram and UpSet plot representation of metabolomics data acquired on cultured Symbiodiniaceae. **(A)** Panel A shows the overlap in detected metabolite features between corals and the cultured Symbiodiniaceae: unique to *M. cavernosa* corals (peach), unique to Symbiodiniaceae (blue) and shared (orange). **(B)** Panel B shows UpSet plot of distribution of metabolite features across three species of cultured Symbiodiniaceae arranged by decreasing frequency.

**Figure 7.**
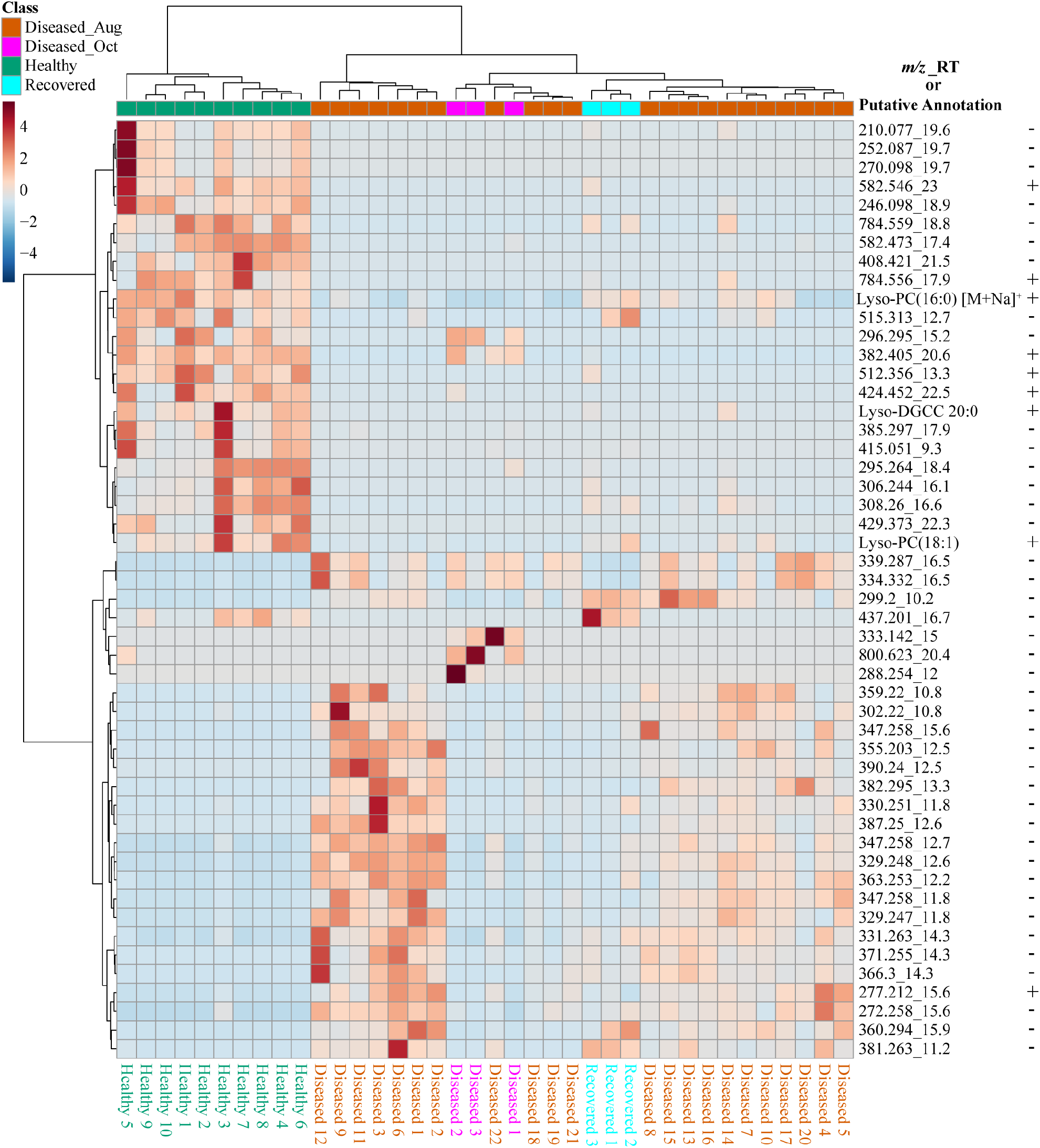
The heat map of top 51 significant features driving the separation of healthy, diseased, and recovered coral samples identified via analysis of variance from LC-MS data using MetaboAnalyst. The scale bar represents autoscaled intensity of features. The putative annotations of these features are provided in Supplementary Table 1. The *m/z*_RT column lists the *m/z* value and retention time of each feature, as well as a few putative annotations. The +/- indicates presence or absence of this feature in the LC-MS data acquired on cultured Symbiodiniaceae.

To gain chemical insights into the overlapping features detected in extracts of coral and cultured Symbiodiniaceae, we attempted annotation of the metabolite features scored in the MDS analysis in Figure 4B, PLS_DA in Supplementary Figure 2 and ANOVA selected features in Figure 7 using FBMN and *in silico* approaches. The features in this category belonged to lipids, including betaine lipids such as diacylglyceryl-carboxyhydroxymethylcholine (e.g., DGCC, *m/z* 490.374), monogalactosyl- and digalactosyldiacylglycerols (MGDG and DGDG), sphingolipids, lyso-PC, steroid-like molecules and tocomonoenol (Supplementary Table 1). Among these, betaine lipids are characterized by the presence of trimethylated hydroxyamino acid connected to the diacylglycerol moiety through an ether bond. To generate a comprehensive list of betaine lipids detected, feature-based molecular networking (Figure 8A) was paired with discovery of mass structural motif specific to betaine lipids using MS2LDA analysis (Figure 8B). The annotation of mass structural motif in Figure 8B allowed annotation of additional betaine lipids not present in the cluster in Figure 8A (and Supplementary Table 2). A comprehensive network of betaine lipids including the ones produced in the cultured Symbiodiniaceae is shown in Supplementary Figure 4. Of note, some ether linked higher chain length acyl tails were observed only in culture extracts of Symbiodiniaceae observed via presence of nodes with higher *m/z* (Supplementary Figure 4). The distribution of a subset of annotated betaine lipids is shown as heat maps in Figure 8C and distribution of all annotated betaine lipids is shown in Supplementary Figure 5. Notably, the healthy sample 3 displayed the largest variation in metabolomics data from PCA plot and also displayed the largest diversity of betaine lipids (Supplementary Figure 5). The betaine lipids have been previously reported in endosymbionts from various coral species (Rosset et al., 2019; Sikorskaya, 2020; Sikorskaya et al., 2021) and are reported as biomarkers of bleaching history (Roach et al., 2021) and thermal tolerance (Leblond et al., 2015; Rosset et al., 2019). It has been previously suggested that betaine lipids being the component of cell membrane are influenced by the host genotype (Sikorskaya et al., 2021). The temporal changes in profiles of these lipids as well as differences between different species undergoing SCTLD in the future will provide important insights into their functions and potential role in resilience.

**Figure 8.**
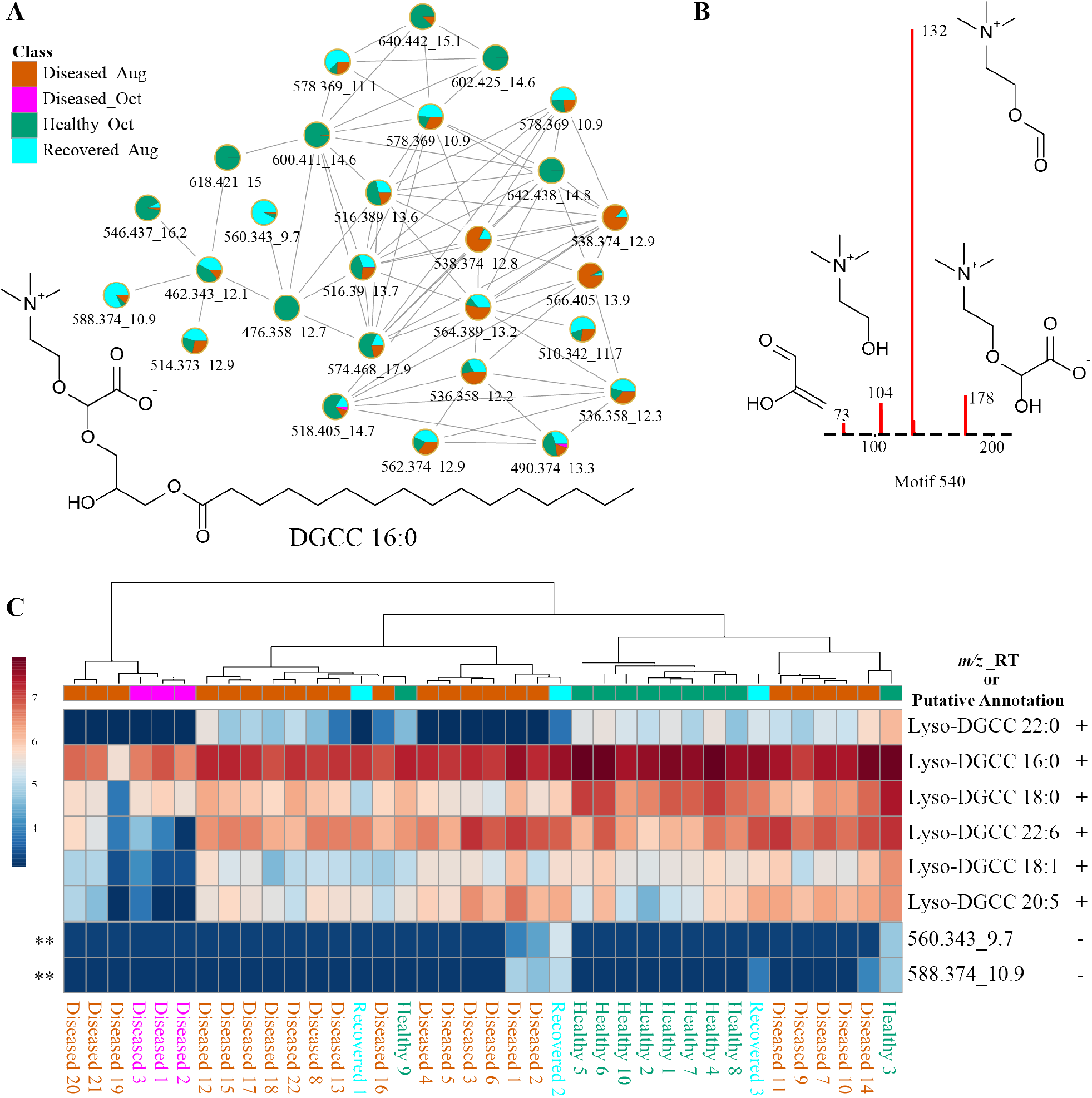
Description of betaine lipids using molecular networking, substructure discovery via MS2LDA, and distribution in corals via heat maps. **(A)** Panel A shows the FBMN of betaine lipids. **(B)** Panel B shows specific fragments observed in the MS^2^ spectra of betaine lipids that constitute the MS2LDA sub-structure motif specific to the head group of betaine lipids. This motif enabled annotation of additional betaine lipids not observed in the network of nodes in panel A. **(C)** Heat map of six betaine lipids across healthy, diseased, and recovered coral samples grouped via analysis of variance from LC-MS generated in MetaboAnalyst. The scale bar represents log-transformed intensity of these lipids. This figure illustrates a subset of betaine lipids. A comprehensive list of all betaine lipids annotated in this study and the heat map for these features is provided in Supplementary Table 2 and Supplementary Figure 5. The *m/z_RT* value or putative annotation are included for each feature. The +/- indicates presence or absence of this feature in the LC-MS data acquired on cultured Symbiodiniaceae.

Glycolipids are present in thylakoid membranes of Symbiodiniaceae and their importance in heat tolerance has been previously reported (Tchernov et al., 2004; Leblond et al., 2015; Rosset et al., 2019). The glycolipid cluster in FBMN (Figure 9A) was annotated using MolDiscovery, which compares user uploaded MS/MS spectra with *in-silico* generated MS/MS spectra of small molecules, and the annotations were propagated using FBMN and MS2LDA and compared to mass spectrometry data in the literature. The feature with *m/z* 797.519 was annotated as MGDG, potentially heterosigma-glycolipid II, a diacylglycerol glycolipid that contains omega-3 polyunsaturated fatty acid residues (Kobayashi et al., 1992). The detection of specific fragments with *m/z* 333.243 and 259.206 in the MS^2^ spectra supports the presence of 18 carbons with 4 desaturations as one of the acyl tail (Figure 9B). Therefore, feature *m/z* 797.519 was annotated as a monogalactosyldiacylglycerol, MGDG (20:5/18:4). Additional MGDG glycolipids were annotated using propagation of FBMN (Supplementary Table 3). The fragments with *m/z* 333.243 and 259.206 resulted in a MS2LDA substructure motif specific to galactosyldiacylglycerols containing an 18:4 acyl tail. Using this chemical information, the features *m/z* 771.504 and 769.488 were annotated as MGDG (18:4/18:4) and MGDG (18:5/18:4), respectively. These Symbiodiniaceae produced glycolipids displayed different patterns than Symbiodiniaceae produced betaine lipids in healthy as well as diseased samples (Figure 9C and Supplementary Figure 6). Indeed, glycolipids and not betaine lipids were shown to be variable when *Cladocopium* C3 was exposed to thermal stress (Rosset et al., 2019). These differences suggest that in addition to the variability in the abundance of the Symbiodiniaceae, their chemotypes are also variable across diseased samples. In recent work by Landsberg and colleagues, histopathological examination of several species of corals with tissue loss attributed to SCTLD revealed disruption in physiology of host and Symbiodiniaceae (Landsberg et al., 2020).The differences in Symbiodiniaceae-specific metabolites further support this finding. In algae and dinoflagellates, MGDG are one of the main classes of lipid that comprise thylakoid membranes (Li-Beisson et al., 2019; Sikorskaya et al., 2021), on which light-dependent photosynthesis occurs, and their functions as antitumor, anti-inflammatory, anticancer, and appetite-suppressing agents have been described from plant sources. Their variability among corals undergoing different diseases should be compared and correlated to Symbiodiniaceae genotypes to link their functions to holobiont health. Since changes in glycolipids in thylakoid membranes have been linked to thermotolerance in corals and in other microalgae (Huang et al., 2017) and increased frequency of mass bleaching events are occurring, deeper understanding of these lipid distributions in different species of corals either resilient or susceptible to disease as well as in culture extracts of isolated symbionts is necessary. Previously, GC-MS based metabolomics did not reveal correlation between symbiont composition and the host metabolite profiles (Matthews et al., 2020), but LC-MS based metabolomics of the entire holobiont in our case, clearly reveals metabolite differences that are attributed to the symbionts via a culture-based strategy.

**Figure 9.**
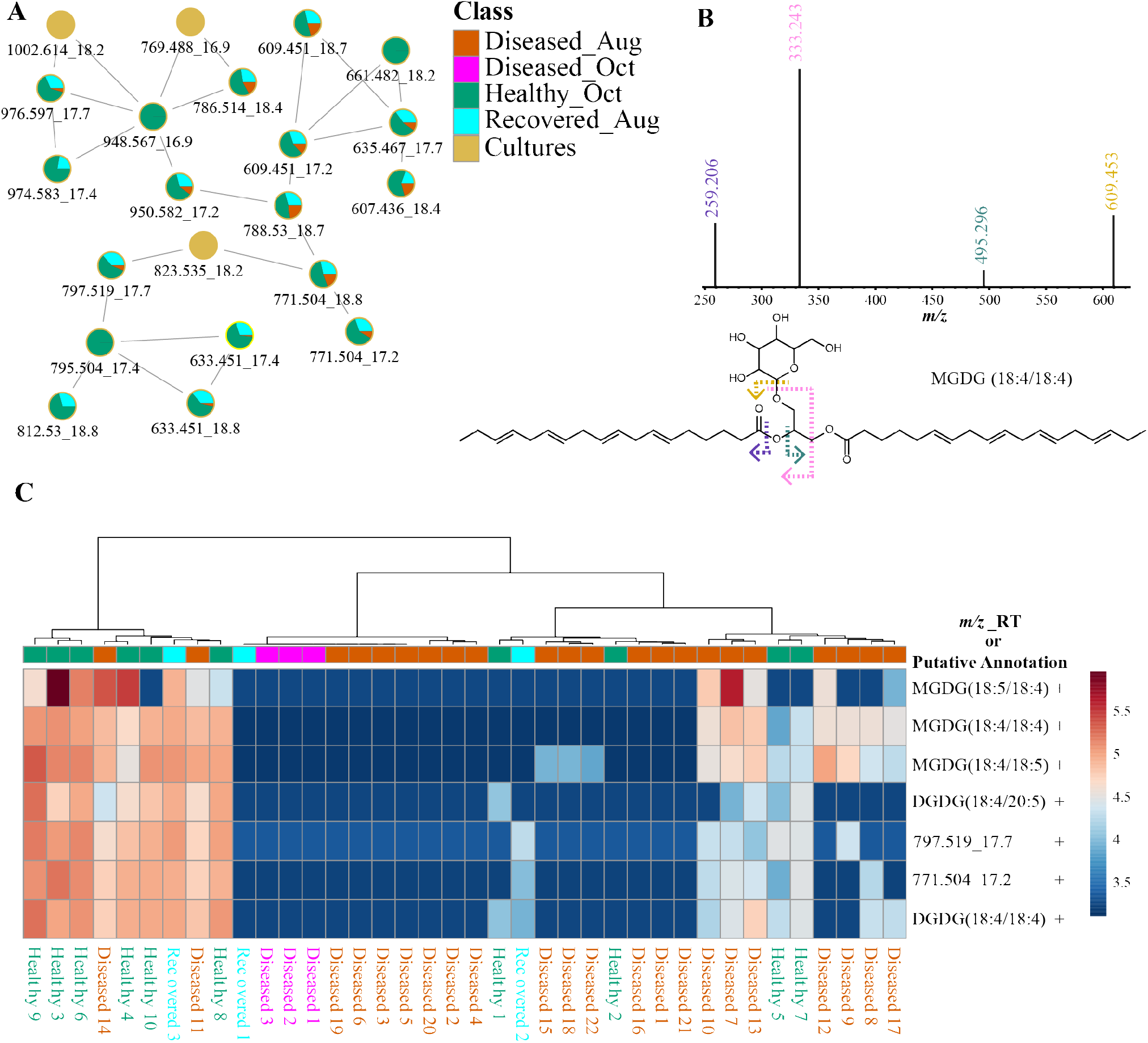
Description of putatively annotated glycolipids using molecular networking, fragmentation spectra, and distribution via heat maps. **(A)** Panel A shows the FBMN of glycolipids. **(B)** Panel B shows specific fragments with *m/z* 333.243 and 259.206 observed in the MS^2^ spectra of glycolipids support the annotation of acyl tail as 18:4. **(C)** Heat map of seven glycolipids across healthy, diseased, and recovered coral samples grouped via analysis of variance from LC-MS data generated using MetaboAnalyst. The scale bar represents log-transformed intensity of these lipids. This figure illustrates a subset of glycolipids. A comprehensive list of all glycolipids and the heat map for these features is provided in Supplementary Table 3 and Supplementary Figure 6. The *m/z*_RT value or putative annotations are included for each feature. The +/- indicates presence or absence of this feature in the LC-MS data acquired on cultured Symbiodiniaceae.

In addition to various lipids such as LysoPC, sphinogolipids, and steroid-like molecules detected at higher levels in healthy corals (Supplementary Table 1), an α-tocopherol analog, tocomonoenol also detected in the Symbiodiniaceae cultures and was among the top ranked features (Figure 10). Detection of tocomonoenol in culture extracts suggests that the origin of this molecule is endosymbiotic Symbiodiniaceae, again supporting that culture-based strategy could facilitate description of individual contributions of each partner of the holobiont. Indeed, the chloroplast-associated antioxidant alpha-tocopherol was observed to be accumulated in intracellular metabolite pools of symbionts during thermal bleaching (Hillyer et al., 2017) and the antioxidant activity of tocomonoenol and other tocopherols have been previously explored (Yamamoto et al., 1999; Beppu et al., 2020). The annotation of the tocomonoenol was enabled by the *in silico* tool MolDiscovery and supported by the MS^2^ spectra for α-tocopherol available in the NIST database (Figure 10B). Thus, in the absence of mass spectral databases, such *in silico* tools serve as a valuable strategy enabling glimpse into structural scaffolds detected in coral holobiont and culture-based strategies allow assignment of biological partner contributing the metabolite (Figure 10). FBMN also allowed visualization of various analogs of α-tocopherol (Figure 10A). The annotation of α-tocopherol was confirmed using commercial standard (Supplementary Figure 7). The mirror plots in Supplementary Figure 8A support that metabolites with *m/z* 424.332 (orange node, Figure 10A) and *m/z* 410.317 (orange and cyan-colored node, Figure 10A) are tocotrienols. While tocopherol with a fully saturated polyprenyl side chain and tocomonoenol with a single unsaturation of the side chain were observed at higher abundances in healthy samples, tocotrienol with three unsaturations of the side chain (*m/z* 410.318) accumulated in diseased corals (Supplementary Figure 8B).

**Figure 10.**
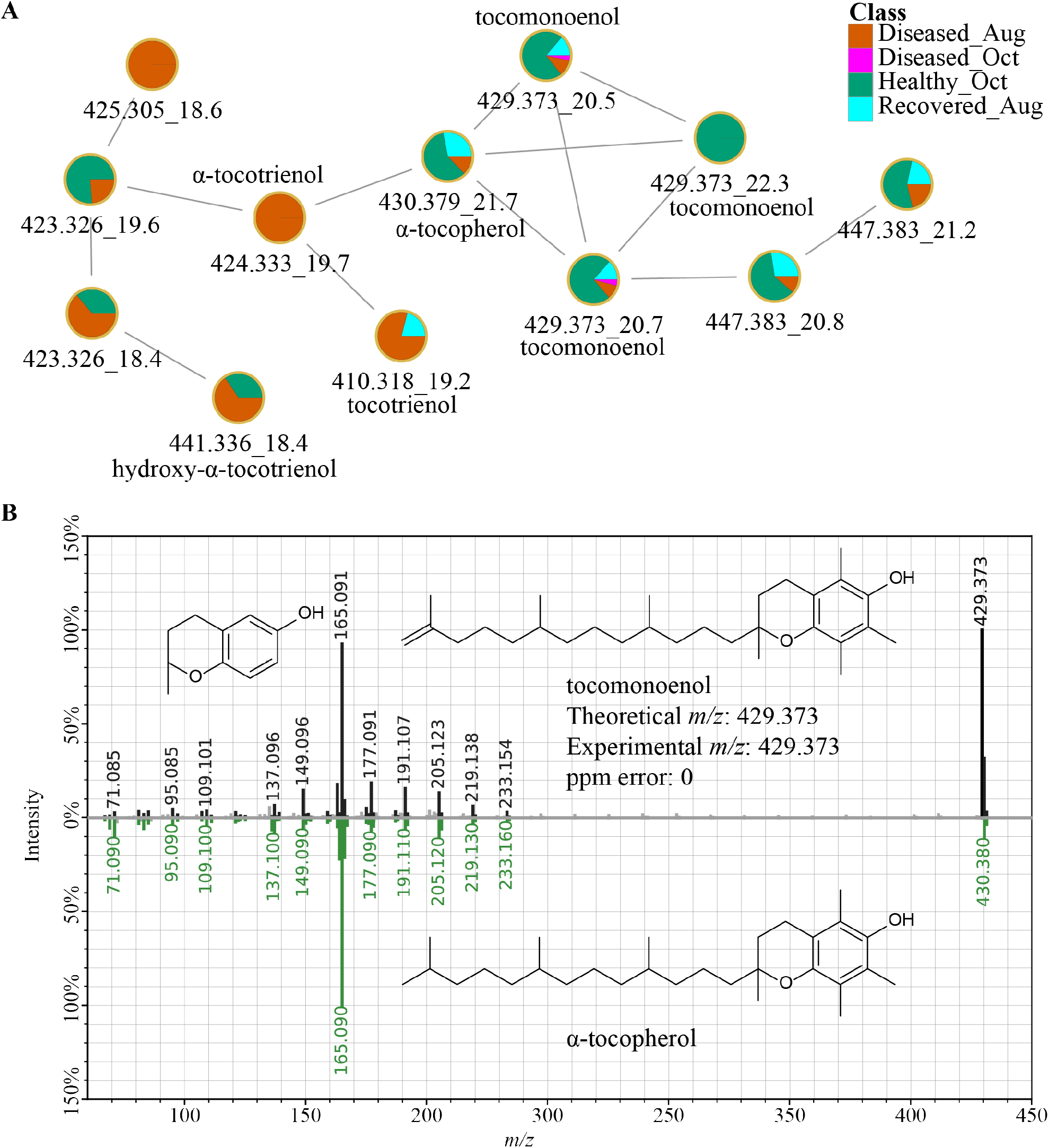
α-Tocopherol and related metabolites. **(A)** Panel A shows the FBMN of tocopherols. The MS^2^-based spectral annotations of these compounds and their distribution in corals analyzed in this study are provided in Supplementary Figures 7 and 8. **(B)** Panel B shows the mirror plot of MS^2^ spectra of tocomonoenol acquired on extracts of coral samples (black trace) and a MS^2^ spectra of tocopherol available in the NIST database (green trace) supporting the annotation of the feature with *m/z* 429.373 as tocomonoenol.

Lastly, the metabolites observed at higher abundances in the diseased samples were queried based on a heat map generated on features ranked via non-parametric ANOVA for log transformed data (Supplementary Figure 9). Several metabolites with similar MS^2^ spectra in this class belonged to the chemical class of unsaturated fatty acids such as docosahexaenoic acid (Supplementary Figure 10 and Supplementary Table 4). Accumulation of unsaturated fatty acids upon stress induced by exposure to high concentrations of pollutant, ethylhexyl salicylate, in the coral *Pocillopora damicornis* has been reported and their potential as biomarker for stress in corals has been suggested (Stien et al., 2020). These observations warrant targeted analysis of polyunsaturated fatty acids and their downstream effectors such as eicosanoids in corals undergoing disease.

## Conclusions

In this study, we employed metabolomic profiling to visualize intraspecies variation of *M. cavernosa* corals with no visual sign of disease, corals that apparently recovered, and corals presumably with SCTLD. The unsupervised PCA, HCA, and MDS revealed metabolomic variation within healthy corals, diseased corals, and between healthy and diseased corals, which was attributed to various metabolites including lipids and tocopherols produced by Symbiodiniaceae. In many instances, the healthy samples that clustered closer in PCA and HCA plots were also immediate neighbors in the site map. Thus, the proximity resulted in similar metabolomes in most cases with a few exceptions. This observation was not true for most diseased colonies, which showed wider spread in metabolomic profiles in the PCA plot. In the future, time series experimental design for disease progression, along with characterization of host genotype, symbiont genotype, and microbiome structure is important to explain the source of this variability in the metabolomic profiles.

The emerging approaches in metabolomics technology allow analyses of hundreds to thousands of metabolites. The major bottleneck in data analyses in metabolomics stems from our inability to annotate the metabolites detected and to assign the biological source of the metabolite to an individual holobiont partner. In the absence of comprehensive mass spectral databases for coral metabolites, annotations were made using *in silico* methods and literatures searches. These compound annotations were propagated to additional unknown metabolite features through a feature based molecular networking approach and by discovery of sub-structural motifs using MS2LDA. Thus, we demonstrate that use of *in silico* tools serve as a practical approach in the absence of databases to aid and improve metabolite identifications for profiling of coral holobionts. Better understanding of holobiont response to disease can be gleaned if metabolomics variation can be further assigned to the respective partner in the holobiont. In this work, we highlight a feature based molecular networking approach to metabolomics data acquired on cultured Symbiodiniaceae and the holobiont to assign sources of detected metabolites for the first time. Using this strategy, we annotated various top features describing metabolomics variation to Symbiodiniaceae demonstrating the usefulness of this approach. A caveat of this culture-based approach is that not all metabolites relevant in the holobiont may be produced in culture. A tedious approach to overcome this limitation is to culture the symbionts under different growth conditions such as temperature variation, nutrient limitation, etc. to induce production of metabolites otherwise observed in a holobiont. In addition, direct extraction of metabolites from the relatively pure fraction of in-hospite Symbiodiniaceae isolated from the coral should be applied for direct comparison. Indeed, this approach has been used previously to describe lipid composition of different species of Symbiodiniaceae under temperature stress. Once the source of a metabolite is assigned to the holobiont partner, genomics can be employed to decipher the biochemical pathways and their regulatory mechanisms. As additional metabolomic datasets become available on cultured coral holobiont partners including cultured bacteria, such analyses will continue to provide biochemical insights into the workings of a coral holobiont. Thus, linking metabolites to their biological producer through comparative metabolomics in a coral holobiont is an important next step in understanding symbiosis in corals and its disruption during disease and environmental stress.

## Author Contributions

JMD conducted metabolomics data acquisition, analyses, data interpretation, and assisted with manuscript editing. OJ assisted with metabolomics data analyses. KP and JH conducted experiments. NG helped with project conception, supervised the project, designed experimental approach, and assisted with various methodological, data and statistical analyses, and wrote the manuscript. VJP helped design the sampling scheme, supervised sample handling and extractions, and contributed to interpretation of data and manuscript editing. BU helped with project conception and editing of manuscript. BW helped with project conception, experimental design, data analyses, and editing of manuscript. All authors contributed to the article and approved the submitted version.

## Acknowledgments

We wish to thank Ricardo R. da Silva (School of Pharmaceutical Sciences of Ribeirão Preto, University of São Paulo, Ribeirão Preto) for discussion on statistical analyses presented in the paper. We thank Kristin Anderson, Sammi Buckley, Brooke Enright, Hunter Noren and Thomas Ingalls for assistance with fieldwork. We thank the Florida Fish and Wildlife Conservation Commission for permitting this activity under the State of Florida Special Activity License Permits SAL-19-1702-SRP,SAL-20-1702-SRP and SAL-19-2201-SRP. We thank Andrew Baker and Richard Karp (University of Miami) for sharing Symbiodiniaceae cultures from the BURR collection provided by Mary Alice Coffroth.

## Data Availability Statement

The LC-MS/MS datasets generated in this study are available in the public repository, Mass spectrometry Interactive Virtual Environment (MassIVE) at https://massive.ucsd.edu/ with the identifier MSV000087471.

## References

Aeby, G.S., Ushijima, B., Campbell, J.E., Jones, S., Williams, G.J., Meyer, J.L., et al. (2019). Pathogenesis of a tissue loss disease affecting multiple species of corals along the Florida reef tract. Front. Mar. Sci. 6, 678. doi: 10.3389/fmars.2019.00678.

Ainsworth, T.D., Krause, L., Bridge, T., Torda, G., Raina, J.-B., Zakrzewski, M., et al. (2015). The coral core microbiome identifies rare bacterial taxa as ubiquitous endosymbionts. ISME Journal 9(10), 2261–2274. doi: 10.1038/ismej.2015.39.

Alvarez-Filip, L., Estrada-Saldívar, N., Pérez-Cervantes, E., Molina-Hernández, A., and González-Barrios, F.J. (2019). A rapid spread of the stony coral tissue loss disease outbreak in the Mexican Caribbean. PeerJ 7, e8069. doi: 10.7717/peerj.8069.

Andersson, E.R., Day, R.D., Loewenstein, J.M., Woodley, C.M., and Schock, T.B. (2019). Evaluation of sample preparation methods for the analysis of reef-building corals using 1H-NMR-based metabolomics. Metabolites 9(2), 32.

Baker, A.C. (2003). Flexibility and specificity in coral-algal symbiosis: diversity, ecology, and biogeography of *Symbiodinium*. Annu. Rev. Ecol. Evol. Syst. 34(1), 661–689. doi: 10.1146/annurev.ecolsys.34.011802.132417.

Becker, C.C., Brandt, M., Miller, C.A., and Apprill, A. (2021). Stony Coral Tissue Loss Disease biomarker bacteria identified in corals and overlying waters using a rapid field-based sequencing approach. bioRxiv, 2021.2002.2017.431614. doi: 10.1101/2021.02.17.431614.

Benkwitt, C.E., Wilson, S.K., and Graham, N.A.J. (2020). Biodiversity increases ecosystem functions despite multiple stressors on coral reefs. Nat. Ecol. Evol. (4), 919–926. doi: 10.1038/s41559-020-1203-9.

Beppu, F., Aida, Y., Kaneko, M., Kasatani, S., Aoki, Y., and Gotoh, N. (2020). Functional evaluation of marine-derived tocopherol, a minor homolog of vitamin E, on adipocyte differentiation and inflammation using 3T3-L1 and RAW264.7 cells. Fish. Sci. 86(2), 415–425. doi: 10.1007/s12562-020-01404-6.

Brander, L.M., and Van Beukering, P. (2013). The total economic value of U.S. coral reefs: a review of the literature. Available at: https://repository.library.noaa.gov/view/noaa/8951.

Bruckner, A.W. (2002). Life-saving products from coral reefs. Issues Sci. Technol. 18(3), 39–44.

Budd, A.F., Nunes, F.L.D., Weil, E., and Pandolfi, J.M. (2012). Polymorphism in a common Atlantic reef coral (*Montastraea cavernosa*) and its long-term evolutionary implications. Evol. Ecol. 26(2), 265–290. doi: 10.1007/s10682-010-9460-8.

Cao, L., Guler, M., Tagirdzhanov, A., Lee, Y., Gurevich, A., and Mohimani, H. (2020). MolDiscovery: Learning mass spectrometry fragmentation of small molecules. bioRxiv, 2020.2011.2028.401943. doi: 10.1101/2020.11.28.401943.

Chong, J., Wishart, D.S., and Xia, J. (2019). Using metaboAnalyst 4.0 for comprehensive and integrative metabolomics data analysis. Curr. Protoc. Bioinformatics 68(1), e86. doi: 10.1002/cpbi.86.

Correa, A.M.S., Brandt, M.E., Smith, T.B., Thornhill, D.J., and Baker, A.C. (2009). *Symbiodinium* associations with diseased and healthy scleractinian corals. Coral Reefs 28(2), 437–448. doi: 10.1007/s00338-008-0464-6.

Dixon, G.B., Davies, S.W., Aglyamova, G.V., Meyer, E., Bay, L.K., and Matz, M.V. (2015). Genomic determinants of coral heat tolerance across latitudes. Science 348(6242), 1460–1462. doi: 10.1126/science.1261224.

Djoumbou, F.Y., Eisner, R., Knox, C., Chepelev, L., Hastings, J., Owen, G., et al. (2016). ClassyFire: automated chemical classification with a comprehensive, computable taxonomy. Journal of Cheminformatics 8(1), 61. doi: 10.1186/s13321-016-0174-y.

Drake, J.L., Schaller, M.F., Mass, T., Godfrey, L., Fu, A., Sherrell, R.M., et al. (2018). Molecular and geochemical perspectives on the influence of CO_2_ on calcification in coral cell cultures. Limnology and Oceanography 63(1), 107–121. doi: 10.1002/lno.10617.

Dührkop, K., Fleischauer, M., Ludwig, M., Aksenov, A.A., Melnik, A.V., Meusel, M., et al. (2019). SIRIUS 4: a rapid tool for turning tandem mass spectra into metabolite structure information. Nat. Methods 16(4), 299–302. doi: 10.1038/s41592-019-0344-8.

Dührkop, K., Nothias, L.-F., Fleischauer, M., Reher, R., Ludwig, M., Hoffmann, M.A., et al. (2021). Systematic classification of unknown metabolites using high-resolution fragmentation mass spectra. Nat. Biotechnol. 39(4), 462–471. doi: 10.1038/s41587-020-0740-8.

Estrada-Saldívar, N., Molina-Hernández, A., Pérez-Cervantes, E., Medellín-Maldonado, F., González-Barrios, F.J., and Alvarez-Filip, L. (2020). Reef-scale impacts of the stony coral tissue loss disease outbreak. Coral Reefs 39(4), 861–866. doi: 10.1007/s00338-020-01949-z.

Estrada-Saldívar, N., Quiroga-García, B.A., Pérez-Cervantes, E., Rivera-Garibay, O.O., and Alvarez-Filip, L. (2021). Effects of the stony coral tissue loss disease outbreak on coral communities and the benthic composition of cozumel reefs. Front. Mar. Sci. 8(306), 632777. doi: 10.3389/fmars.2021.632777.

Fisher, R., O’Leary, Rebecca A., Low-Choy, S., Mengersen, K., Knowlton, N., Brainard, Russell E., et al. (2015). Species richness on coral reefs and the pursuit of convergent global estimates. Curr. Biol. 25(4), 500–505. doi: 10.1016/j.cub.2014.12.022.

Garg, N., Wang, M., Hyde, E., da Silva, R.R., Melnik, A.V., Protsyuk, I., et al. (2017). Three-dimensional microbiome and metabolome cartography of a diseased human lung. Cell Host Microbe 22(5), 705–716. doi: 10.1016/j.chom.2017.10.001.

Goodacre, R. (2007). Metabolomics of a superorganism. J. Nutr. 137(1), 259S–266S. doi: 10.1093/jn/137.1.259S.

Gordon, B.R., and Leggat, W. (2010). *Symbiodinium*-invertebrate symbioses and the role of metabolomics. Mar. Drugs. 8(10), 2546–2568. doi: 10.3390/md8102546.

Gordon, B.R., Leggat, W., and Motti, C.A. (2013). “Extraction protocol for nontargeted NMR and LC-MS metabolomics-based analysis of hard coral and their algal symbionts,” in Metabolomics tools for natural product discovery. Springer, 129–147.

Gowda, H., Ivanisevic, J., Johnson, C.H., Kurczy, M.E., Benton, H.P., Rinehart, D., et al. (2014). Interactive XCMS Online: simplifying advanced metabolomic data processing and subsequent statistical analyses. Anal. Chem. 86(14), 6931–6939. doi: 10.1021/ac500734c.

Grottoli, A.A.-O., Dalcin Martins, P., Wilkins, M.J., Johnston, M.D., Warner, M.E., Cai, W.J., et al. (2018). Coral physiology and microbiome dynamics under combined warming and ocean acidification. PLOS One 13(1), e0191156. doi: 10.1371/journal.pone.0191156.

Heres, M.M., Farmer, B.H., Elmer, F., and Hertler, H. (2021). Ecological consequences of stony coral tissue loss disease in the Turks and Caicos Islands. Coral Reefs 40(2), 609–624. doi: 10.1007/s00338-021-02071-4.

Hillyer, K.E., Dias, D.A., Lutz, A., Wilkinson, S.P., Roessner, U., and Davy, S.K. (2017). Metabolite profiling of symbiont and host during thermal stress and bleaching in the coral *Acropora aspera*. Coral Reefs 36(1), 105–118. doi: 10.1007/s00338-016-1508-y.

Hoegh-Guldberg, O., Poloczanska, E.S., Skirving, W., and Dove, S. (2017). Coral reef ecosystems under climate change and ocean acidification. Front. Mar. Sci. 4, 158. doi: 10.3389/fmars.2017.00158.

Howells, E.J., Vaughan, G.O., Work, T.M., Burt, J.A., and Abrego, D. (2020). Annual outbreaks of coral disease coincide with extreme seasonal warming. Coral Reefs 39(3), 771–781. doi: 10.1007/s00338-020-01946-2.

Huang, L., Xu, J., Zong, C., Zhu, S., Ye, M., Zhou, C., et al. (2017). Effect of high temperature on the lipid composition of *Isochrysis galbana* Parke in logarithmic phase. Aquac. Int. 25(1), 327–339. doi: 10.1007/s10499-016-0031-z.

Joyce, A.R., and Palsson, B.Ø. (2006). The model organism as a system: integrating ‘omics’ data sets. Nat. Rev. Mol. Cell Biol. 7(3), 198–210. doi: 10.1038/nrm1857.

Kenkel, C.D., and Matz, M.V. (2016). Gene expression plasticity as a mechanism of coral adaptation to a variable environment. Nat. Ecol. Evol. 1(1), 0014. doi: 10.1038/s41559-016-0014.

Kobayashi, M., Hayashi, K., Kawazoe, K., and Kitagawa, I. (1992). Marine natural products. XXIX. *Heterosigma*-glycolipids I, II, III, and IV, four diacylglyceroglycolipids possessing ω3-polyunsaturated fatty acid residues, from the raphidopycean dinoflagellate *Heterosigma akashiwo*. Chem. Pharm. Bull. 40(6), 1404–1410.

Kramer, P.R., Roth, L., and Lang, J. (2020). Map of Coral Cover of Susceptible Coral Species to SCTLD. www.agrra.org. ArcGIS Online.

LaJeunesse, T.C., Parkinson, J.E., Gabrielson, P.W., Jeong, H.J., Reimer, J.D., Voolstra, C.R., et al. (2018). Systematic revision of Symbiodiniaceae highlights the antiquity and diversity of coral endosymbionts. Curr. Biol. 28(16), 2570–2580. doi: 10.1016/j.cub.2018.07.008.

Landsberg, J.H., Kiryu, Y., Peters, E.C., Wilson, P.W., Perry, N., Waters, Y., et al. (2020). Stony coral tissue loss disease in Florida is associated with disruption of host-zooxanthellae physiology. Front. Mar. Sci. 7, 1090. doi: doi.org/10.3389/fmars.2020.576013.

Leblond, J.D., Khadka, M., Duong, L., and Dahmen, J.L. (2015). Squishy lipids: Temperature effects on the betaine and galactolipid profiles of a C18/C18 peridinin-containing dinoflagellate, *Symbiodinium microadriaticum* (Dinophyceae), isolated from the mangrove jellyfish, *Cassiopea xamachana*. Phycol. Res. 63(3), 219–230. doi: 10.1111/pre.12093.

Li-Beisson, Y., Thelen, J.J., Fedosejevs, E., and Harwood, J.L. (2019). The lipid biochemistry of eukaryotic algae. Prog. Lipid Res. 74, 31–68. doi: 10.1016/j.plipres.2019.01.003.

Littman, R.A., Bourne, D.G., and Willi, B.L. (2010). Responses of coral-associated bacterial communities to heat stress differ with *Symbiodinium* type on the same coral host. Mol. Ecol. 19(9), 1978–1990. doi: 10.1111/j.1365-294X.2010.04620.x.

Lohr, K.E., Khattri, R.B., Guingab-Cagmat, J., Camp, E.F., Merritt, M.E., Garrett, T.J., et al. (2019). Metabolomic profiles differ among unique genotypes of a threatened Caribbean coral. Sci. Rep 9(1), 6067. doi: 10.1038/s41598-019-42434-0.

Maire, J., Girvan, S.K., Barkla, S.E., Perez-Gonzalez, A., Suggett, D.J., Blackall, L.L., et al. (2021). Intracellular bacteria are common and taxonomically diverse in cultured and in hospite algal endosymbionts of coral reefs. ISME J. doi: 10.1038/s41396-021-00902-4.

Maruyama, S., and Weis, V.M. (2021). Limitations of using cultured algae to study cnidarian-algal symbioses and suggestions for future studies. J. Phycol. 57(1), 30–38. doi: 10.1111/jpy.13102.

Matthews, J.L., Cunning, R., Ritson-Williams, R., Oakley, C.A., Lutz, A., Roessner, U., et al. (2020). Metabolite pools of the reef building coral *Montipora capitata* are unaffected by Symbiodiniaceae community composition. Coral Reefs 39(6), 1727–1737. doi: 10.1007/s00338-020-01999-3.

Maynard, J., van Hooidonk, R., Eakin, C.M., Puotinen, M., Garren, M., Williams, G., et al. (2015). Projections of climate conditions that increase coral disease susceptibility and pathogen abundance and virulence. Nat. Clim. Change. 5(7), 688–694. doi: 10.1038/nclimate2625.

McAvoy, A.C., Jaiyesimi, O., Threatt, P.H., Seladi, T., Goldberg, J.B., da Silva, R.R., et al. (2020). Differences in cystic fibrosis-associated *Burkholderia* spp. bacteria metabolomes after exposure to the antibiotic trimethoprim. ACS Infect. Dis. 6(5), 1154–1168. doi: 10.1021/acsinfecdis.9b00513.

Meiling, S.S., Muller, E.M., Lasseigne, D., Rossin, A., Veglia, A.J., MacKnight, N., et al. (2021). Variable species responses to experimental stony coral tissue loss disease (SCTLD) exposure. Front. Mar. Sci. 8(464), 670829. doi: 10.3389/fmars.2021.670829.

Melnik, A.V., Vazquez-Baeza, Y., Aksenov, A.A., Hyde, E., McAvoy, A.C., Wang, M., et al. (2019). Molecular and microbial microenvironments in chronically diseased lungs associated with cystic fibrosis. mSystems 4(5), e00375–00319. doi: 10.1128/mSystems.00375-19.

Mera, H., and Bourne, D.G. (2018). Disentangling causation: complex roles of coral-associated microorganisms in disease. Environ. Microbiol. 20(2), 431–449. doi: 10.1111/1462-2920.13958.

Meyer, J.L., Castellanos-Gell, J., Aeby, G.S., Häse, C.C., Ushijima, B., and Paul, V.J. (2019). Microbial community shifts associated with the ongoing stony coral tissue loss disease outbreak on the Florida reef tract. Front. Microbiol. 10, 2244. doi: 10.3389/fmicb.2019.02244.

Micallef, L., and Rodgers, P. (2014). eulerAPE: Drawing Area-Proportional 3-Venn Diagrams Using Ellipses. PLOS One 9(7), e101717.. doi: 10.1371/journal.pone.0101717.

Mieog, J.C., Olsen, J.L., Berkelmans, R., Bleuler-Martinez, S.A., Willis, B.L., and van Oppen, M.J.H. (2009). The roles and interactions of symbiont, host and environment in defining coral fitness. PLoS One 4(7), e6364. doi: 10.1371/journal.pone.0006364.

Miller, M.W., Karazsia, J., Groves, C.E., Griffin, S., Moore, T., Wilber, P., et al. (2016). Detecting sedimentation impacts to coral reefs resulting from dredging the Port of Miami, Florida USA. Peer J 4, e2711.

Montilla, L.M., Ascanio, A., Verde, A., and Croquer, A. (2019). Systematic review and meta-analysis of 50 years of coral disease research visualized through the scope of network theory. PeerJ 7, e7041. doi: 10.7717/peerj.7041.

Muller, E.M., Sartor, C., Alcaraz, N.I., and van Woesik, R. (2020). Spatial epidemiology of the Stony-Coral-Tissue-Loss disease in Florida. Front. Mar. Sci. 7, 163. doi: 10.3389/fmars.2020.00163.

Muscatine, L. (1990). The role of symbiotic algae in carbon and energy flux in reef corals. Coral Reefs 25, 75–87.

Nakamura, H., Kawase, Y., Maruyama, K., and Murai, A. (1998). Studies on polyketide metabolites of a symbiotic dinoflagellate, *Symbiodinium* sp.: A new C30 marine alkaloid zooxanthellamine, a plausible precursor for zoanthid alkaloids. Bull. Chem. Soc. Jpn. 71(4), 781–787.

Neely, K.L., Macaulay, K.A., Hower, E.K., and Dobler, M.A. (2020). Effectiveness of topical antibiotics in treating corals affected by Stony Coral Tissue Loss Disease. PeerJ 8, e9289. doi: 10.7717/peerj.9289.

Nothias, L.-F., Petras, D., Schmid, R., Dührkop, K., Rainer, J., Sarvepalli, A., et al. (2020). Feature-based molecular networking in the GNPS analysis environment. Nat. Methods 17(9), 905–908. doi: 10.1038/s41592-020-0933-6.

Ochsenkühn, M.A., Schmitt-Kopplin, P., Harir, M., and Amin, S.A. (2018). Coral metabolite gradients affect microbial community structures and act as a disease cue. Commun. Biol. 1(1), 184. doi: 10.1038/s42003-018-0189-1.

Pandolfi, J.M., Connolly, S.R., Marshall, D.J., and Cohen, A.L. (2011). Projecting coral reef futures under global warming and ocean acidification. Science 333(6041), 418–422. doi: 10.1126/science.1204794.

Parkinson, J.E., Banaszak, A.T., Altman, N.S., LaJeunesse, T.C., and Baums, I.B. (2015). Intraspecific diversity among partners drives functional variation in coral symbioses. Sci. Rep. 5(1), 15667. doi: 10.1038/srep15667.

Patti, G.J., Yanes, O., and Siuzdak, G. (2012). Metabolomics: the apogee of the omics trilogy. Nat. Rev. Mol. Cell Biol. 13(4), 263–269. doi: 10.1038/nrm3314.

Plaisance, L., Caley, M.J., Brainard, R.E., and Knowlton, N. (2011). The diversity of coral reefs: what are we missing? PLoS One 6(10), e25026. doi: 10.1371/journal.pone.0025026.

Pluskal, T., Castillo, S., Villar-Briones, A., and Orešič, M. (2010). MZmine 2: Modular framework for processing, visualizing, and analyzing mass spectrometry-based molecular profile data. BMC Bioinformatics 11(1), 395. doi: 10.1186/1471-2105-11-395.

Precht, W.F., Gintert, B.E., Robbart, M.L., Fura, R., and van Woesik, R. (2016). Unprecedented disease-related coral mortality in southeastern Florida. Sci. Rep. 6, 31374. doi: 10.1038/srep31374.

Quinn, R.A., Phelan, V.V., Whiteson, K.L., Garg, N., Bailey, B.A., Lim, Y.W., et al. (2016a). Microbial, host and xenobiotic diversity in the cystic fibrosis sputum metabolome. ISME J 10(6), 1483–1498. doi: 10.1038/ismej.2015.207.

Quinn, R.A., Vermeij, M.J.A., Hartmann, A.C., Galtier d’Auriac, I., Benler, S., Haas, A., et al. (2016b). Metabolomics of reef benthic interactions reveals a bioactive lipid involved in coral defence. Proc. R. Soc. B. 283(1829), 20160469. doi: 10.1098/rspb.2016.0469.

Ramos-Silva, P., Kaandorp, J., Huisman, L., Marie, B., Zanella-Cléon, I., Guichard, N., et al. (2013). The skeletal proteome of the coral *Acropora millepora:* The evolution of calcification by co-option and domain shuffling. Mol. Biol. Evol. 30(9), 2099–2112. doi: 10.1093/molbev/mst109.

Reguero, B.G., Storlazzi, C.D., Gibbs, A.E., Shope, J.B., Cole, A.D., Cumming, K.A., et al. (2021). The value of US coral reefs for flood risk reduction. Nat. Sustain. doi: 10.1038/s41893-021-00706-6.

Richardson, L.L. (1998). Coral diseases: what is really known? Trends in ecology & evolution 13(11), 438–443. doi: 10.1016/s0169-5347(98)01460-8.

Roach, T.N.F., Dilworth, J., H, C.M., Jones, A.D., Quinn, R.A., and Drury, C. (2021). Metabolomic signatures of coral bleaching history. Nat. Ecol. Evol. doi: 10.1038/s41559-020-01388-7.

Roach, T.N.F., Little, M., Arts, M.G.I., Huckeba, J., Haas, A.F., George, E.E., et al. (2020). A multiomic analysis of in situ coral-turf algal interactions. Proc. Natl. Acad. Sci. U.S.A. 117(24), 13588–13595. doi: 10.1073/pnas.1915455117%J Proceedings of the National Academy of Sciences.

Rogers, C.S., and Weil, E. (2010). “Coral reef diseases in the Atlantic-Caribbean,” in Coral reefs: an ecosystem in transition, eds. D. Zvy & S. Noga. Springer, 465–492.

Rohwer, F., Seguritan, V., Azam, F., and Knowlton, N. (2002). Diversity and distribution of coral-associated bacteria. Mar. Ecol. Prog. Ser. 243, 1–10.

Rosales, S.M., Clark, A.S., Huebner, L.K., Ruzicka, R.R., and Muller, E.M. (2020). Rhodobacterales and Rhizobiales are associated with Stony Coral Tissue Loss Disease and its suspected sources of transmission. Front. Microbiol. 11, 681. doi: 10.3389/fmicb.2020.00681.

Rosenberg, E., and Ben-Haim, Y. (2002). Microbial diseases of corals and global warming. Environ. Microbiol. 4(6), 318–326. doi: 10.1046/j.1462-2920.2002.00302.x.

Rosset, S., Koster, G., Brandsma, J., Hunt, A.N., Postle, A.D., and D’Angelo, C. (2019). Lipidome analysis of Symbiodiniaceae reveals possible mechanisms of heat stress tolerance in reef coral symbionts. Coral Reefs 38(6), 1241–1253. doi: 10.1007/s00338-019-01865-x.

Roth, M.S. (2014). The engine of the reef: photobiology of the coral-algal symbiosis. Front. Microbiol. 5, 422. doi: 10.3389/fmicb.2014.00422.

Rouzé, H., Lecellier, G., Saulnier, D., and Berteaux-Lecellier, V. (2016). Symbiodinium clades A and D differentially predispose *Acropora cytherea* to disease and *Vibrio* spp. colonization. Ecol. Evol. 6(2), 560–572. doi: 10.1002/ece3.1895.

Sang, V.T., Dat, T.T., Vinh, L.B., Cuong, L.C., Oanh, P.T., Ha, H., et al. (2019). Coral and Coral-Associated Microorganisms: A Prolific Source of Potential Bioactive Natural Products. Mar. Drugs 17(8), 468. doi: 10.3390/md17080468.

Shannon, P., Markiel, A., Ozier, O., Baliga, N.S., Wang, J.T., Ramage, D., et al. (2003). Cytoscape: a software environment for integrated models of biomolecular interaction networks. Genome Res. 13(11), 2498–2504. doi: 10.1101/gr.1239303.

Shilling, E.N., Combs, I.R., and Voss, J.D. (2021). Assessing the effectiveness of two intervention methods for stony coral tissue loss disease on *Montastraea cavernosa*. Sci. Rep. 11(1), 8566. doi: 10.1038/s41598-021-86926-4.

Sikorskaya, T.V. (2020). Investigation of the total lipidome from a *Zoantharia palythoa* sp. Chem. Nat. Compd. 56(1), 44–49. doi: 10.1007/s10600-020-02940-4.

Sikorskaya, T.V., Efimova, K.V., and Imbs, A.B. (2021). Lipidomes of phylogenetically different symbiotic dinoflagellates of corals. Phytochemistry 181, 112579. doi: 10.1016/j.phytochem.2020.112579.

Sikorskaya, T.V., and Imbs, A.B. (2020). Coral lipidomes and their changes during coral bleaching. Russ. J. Bioorganic Chem. 46(5), 643–656. doi: 10.1134/S1068162020050234.

Sogin, E.M., Anderson, P., Williams, P., Chen, C.-S., and Gates, R.D. (2014). Application of 1H-NMR metabolomic profiling for reef-building corals. PLoS One 9(10), e111274. doi: 10.1371/journal.pone.0111274.

Sogin, E.M., Putnam, H.M., Anderson, P.E., and Gates, R.D. (2016). Metabolomic signatures of increases in temperature and ocean acidification from the reef-building coral, *Pocillopora damicornis*. Metabolomics 12(4), 71. doi: 10.1007/s11306-016-0987-8.

Sogin, E.M., Putnam, H.M., Nelson, C.E., Anderson, P., and Gates, R.D. (2017). Correspondence of coral holobiont metabolome with symbiotic bacteria, archaea and *Symbiodinium* communities. Environ. Microbiol. Rep. 9(3), 310–315. doi: 10.1111/1758-2229.12541.

Stien, D., Suzuki, M., Rodrigues, A.M.S., Yvin, M., Clergeaud, F., Thorel, E., et al. (2020). A unique approach to monitor stress in coral exposed to emerging pollutants. Sci. Rep. 10(1), 9601. doi: 10.1038/s41598-020-66117-3.

Sumner, L.W., Amberg, A., Barrett, D., Beale, M.H., Beger, R., Daykin, C.A., et al. (2007). Proposed minimum reporting standards for chemical analysis Chemical Analysis Working Group (CAWG) Metabolomics Standards Initiative (MSI). Metabolomics 3(3), 211–221. doi: 10.1007/s11306-007-0082-2.

Tchernov, D., Gorbunov, M.Y., de Vargas, C., Narayan Yadav, S., Milligan, A.J., Häggblom, M., et al. (2004). Membrane lipids of symbiotic algae are diagnostic of sensitivity to thermal bleaching in corals. Proc. Natl. Acad. Sci. U.S.A. 101(37), 13531–13535. doi: 10.1073/pnas.0402907101.

Thome, P.E., Rivera-Ortega, J., Rodríguez-Villalobos, J.C., Cerqueda-García, D., Guzmán-Urieta, E.O., García-Maldonado, J.Q., et al. (2021). Local dynamics of a white syndrome outbreak and changes in the microbial community associated with colonies of the scleractinian brain coral *Pseudodiploria strigosa*. PeerJ 9, e10695. doi: 10.7717/peerj.10695.

Ushijima, B., Meyer, J.L., Thompson, S., Pitts, K., Marusich, M.F., Tittl, J., et al. (2020). Disease Diagnostics and Potential Coinfections by *Vibrio coralliilyticus* During an Ongoing Coral Disease Outbreak in Florida. Frontiers in Microbiology 11, 2682. doi: 10.3389/fmicb.2020.569354.

van der Hooft, J.J.J., Wandy, J., Young, F., Padmanabhan, S., Gerasimidis, K., Burgess, K.E.V., et al. (2017). Unsupervised discovery and comparison of structural families across multiple samples in untargeted metabolomics. Anal. Chem. 89(14), 7569–7577. doi: 10.1021/acs.analchem.7b01391.

Vega Thurber, R., Mydlarz, L.D., Brandt, M., Harvell, D., Weil, E., Raymundo, L., et al. (2020). Deciphering coral disease dynamics: integrating host, microbiome, and the changing environment. Front. Ecol. Evol. 8(402), 575927. doi: 10.3389/fevo.2020.575927.

Viant, M.R. (2008). Recent developments in environmental metabolomics. Mol. Biosyst. 4(10), 980–986. doi: 10.1039/B805354E.

Walker, B.K. (2012). Spatial analyses of benthic habitats to define coral reef ecosystem regions and potential biogeographic boundaries along a latitudinal gradient. PLoS ONE 7(1), e30466. doi: 10.1371/journal.pone.0030466.

Walker, B.K. (2018). Southeast Florida reef-wide Post-Irma coral disease surveys. Florida DEP. Miami, FL. Pp. 1–37.

Walker, B.K., Noren, H., A., B., and Buckley, S. (2020). SE FL ECA reef-building-coral disease intervention and preparation for restoration: Final Report. Florida DEP. Miami, FL., 80p.

Walker, B.K., Riegl, B., and Dodge, R.E. (2008). Mapping coral reef habitats in southeast Florida using a combined technique approach. J. Coast. Res. 24(5 (245)), 1138–1150. doi: 10.2112/06-0809.1.

Walker, B.K., Turner, N.R., Noren, H.K.G., Buckley, S.F., and Pitts, K.A. (2021). Optimizing Stony Coral Tissue Loss Disease (SCTLD) Intervention Treatments on *Montastraea cavernosa* in an Endemic Zone. Front. Mar. Sci. 8(746). doi: 10.3389/fmars.2021.666224.

Walton, C.J., Hayes, N.K., and Gilliam, D.S. (2018). Impacts of a regional, multi-year, multi-species coral disease outbreak in southeast Florida. Front. Mar. Sci. 5, 323. doi: 10.3389/fmars.2018.00323.

Wang, M., Carver, J.J., Phelan, V.V., Sanchez, L.M., Garg, N., Peng, Y., et al. (2016). Sharing and community curation of mass spectrometry data with Global Natural Products Social molecular networking. Nat. Biotechnol. 34(8), 828–837. doi: 10.1038/nbt.3597.

Williams, A., Chiles, E.N., Conetta, D., Pathmanathan, J.S., Cleves, P.A., Putnam, H.M., et al. (2021). Metabolomic shifts associated with heat stress in coral holobionts. Sci. Adv. 7(1), eabd4210. doi: 10.1126/sciadv.abd4210.

Wishart, D.S. (2019). Metabolomics for investigating physiological and pathophysiological processes. Physiol. Rev. 99(4), 1819–1875. doi: 10.1152/physrev.00035.2018.

Yamamoto, Y., Maita, N., Fujisawa, A., Takashima, J., Ishii, Y., and Dunlap, W.C. (1999). A new vitamin E (α-Tocomonoenol) from eggs of the pacific salmon *Oncorhynchus keta*. J. Nat. Prod. 62(12), 1685–1687. doi: 10.1021/np990230v.

